# Identification of Omaveloxolone as An Endoplasmic Reticulum Associated Degradation Inhibitor That Induces Early Apoptotic Signaling in Multiple Myeloma

**DOI:** 10.1101/2025.04.02.646787

**Authors:** Erin M. Kropp, Sho Matono, Olivia Y. Wang, Aaron M. Robida, Malathi Kandarpa, Jineigh L. Grant, Bryndon J. Oleson, Andrew Alt, Moshe Talpaz, Matthew Pianko, Qing Li

## Abstract

Endoplasmic reticulum-associated degradation (ERAD) is essential for maintaining protein homeostasis, yet its regulatory mechanisms remain poorly understood. A major challenge in studying ERAD is the lack of specific inhibitors targeting the ERAD complex. To address this, we conducted a cell-based high-throughput screen using the FDA-repurposing library and identified omaveloxolone (RTA408) as a potent ERAD inhibitor that selectively impairs the degradation of ER luminal and membrane substrates. Beyond its utility in identifying ERAD substrates, RTA408 exhibits strong cytotoxic effects in multiple myeloma (MM), an incurable plasma cell malignancy. RTA408 inhibits ERAD activity and rapidly induces apoptotic signaling via caspase 8 and the death-inducing signaling complex (DISC). Notably, RTA408 is cytotoxic to malignant plasma cells, including those resistant to proteasome inhibitors, and demonstrates in vivo anti-myeloma activity. Our findings establish ERAD inhibitors as valuable tools for dissecting ERAD regulation while also highlighting their potential as therapeutic agents for MM.

**Graphical Abstract:** 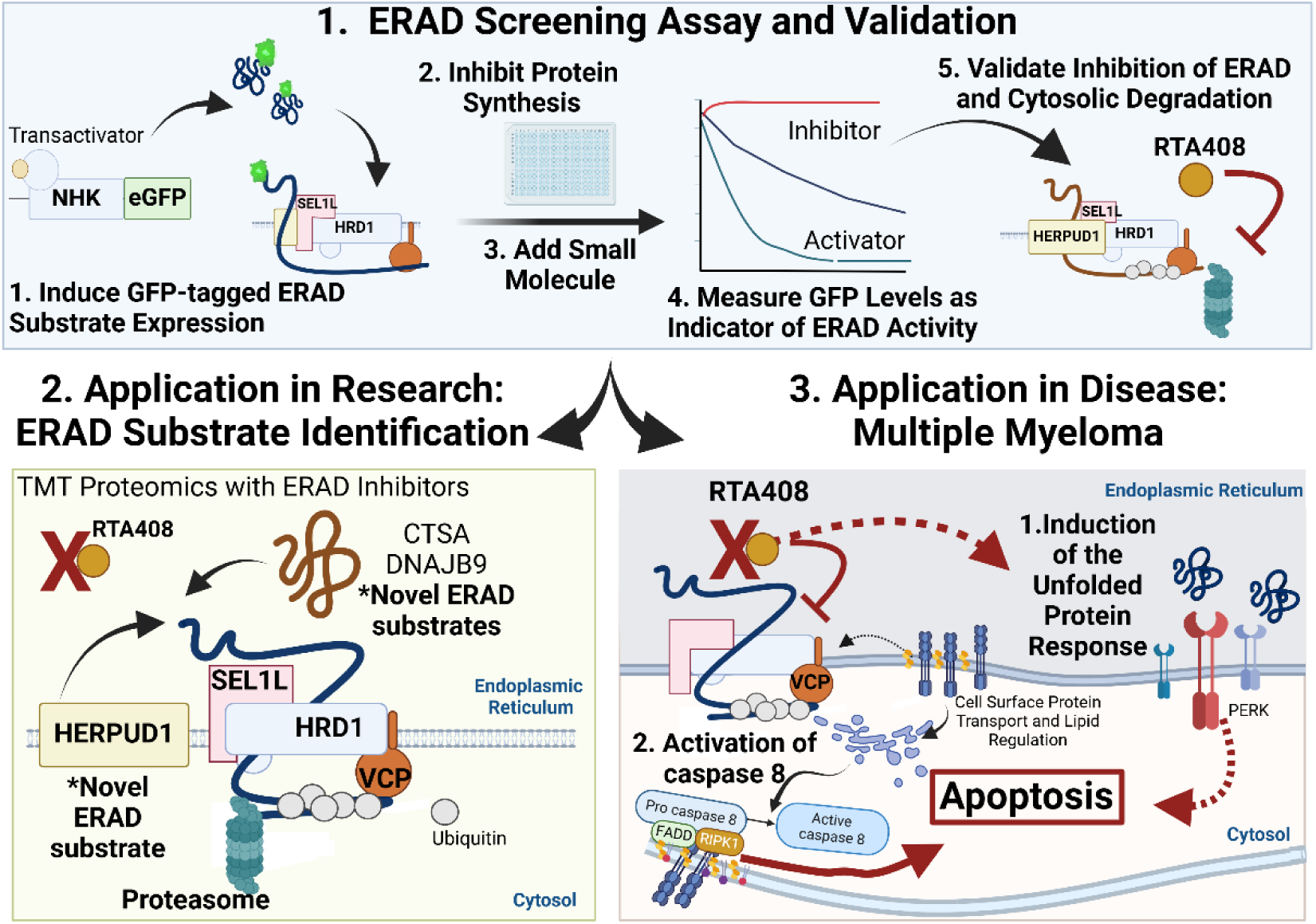

**Highlights:**

- We developed a cell-based screening approach to identify novel modulators of endoplasmic reticulum associated degradation (ERAD), with implications in studying ERAD biology and targeting plasma cell neoplasms.
- Screening of the FDA-repurposing library identified Omaveloxolone (RTA408) as an inhibitor of luminal and membrane ERAD substrate degradation, which can be leveraged to identify ERAD substrates.
- maveloxolone treatment rapidly induces the unfolded protein response and apoptosis that is dependent on caspase 8 and death-inducing signaling complex (DISC) in multiple myeloma cells.
- maveloxolone exhibits cytotoxic effects against multiple myeloma cells *in vitro* and *in vivo* and induces apoptosis in primary plasma cells from patients with relapsed/refractory myeloma

## Introduction

Synthesis of secreted and membrane proteins is a complex process in the endoplasmic reticulum (ER), which is subject to inherent misfolding and stress. Endoplasmic reticulum-associated degradation (ERAD), a crucial component of the ER protein quality control system, is responsible for degrading misfolded ER proteins and regulating ER homeostasis^1^. ERAD is a multiprotein complex that functions to recognize, polyubiquitinate, and translocate target proteins to the cytosol for degradation by the ubiquitin-proteasome system. While ERAD remains an active area of research, its regulatory mechanisms in mammalian cells are not well understood and limited chemical modulators have been identified. Most commercially available compounds do not directly target the ERAD complex but instead inhibit downstream recruitment proteins such as VCP/p97 or the proteasome, which are not specific to ERAD^2–4^. The identification of additional ERAD inhibitors has been hampered by in vitro screening approaches, off-target toxicities, or require high concentrations to inhibit ERAD activity in cells, highlighting the need for novel screening approaches^5,6^.

ERAD has not only been implicated in diseases of protein misfolding or endoplasmic reticulum stress but may be preferentially required in cells with high protein biogenesis, including multiple myeloma (MM). MM is an incurable plasma cell neoplasm characterized by high paraprotein secretion in over 95% of cases^7,8^. Paraproteins are monoclonal immunoglobulins that are synthesized and post-translationally modified in the ER. Immunoglobulin synthesis is subject to inherent protein misfolding, which if unabated, can lead to ER stress and cell death^9,10^. In MM, ERAD disruption has been associated with an induction of ER stress, activation of the unfolded protein response (UPR) pathway, and induction of apoptosis in MM cells^7,9,11^. Furthermore, several ERAD complex proteins are essential for MM cell survival but dispensable in most other cancerous and healthy cell lines, suggesting a unique reliance of MM cells on ERAD activity ^12^. Proteasome inhibitors (PIs) are a cornerstone of MM treatment, disrupting protein degradation via the ubiquitin-proteasome system. PIs inhibit ERAD substrate degradation, leading to ER stress and pro-apoptotic signaling ^9,11^. While PIs have contributed to an improvement in MM overall survival from 2-3 years to 6.5-10 years, almost all patients develop PI resistance resulting in relapsed or refractory (R/R) disease^13,14^. Targeting alternative ERAD proteins has been proposed as a therapeutic strategy to overcome resistance^2,4^. Previous studies have shown that inhibiting specific ERAD proteins is cytotoxic to MM cells and can bypass PI resistance. However, clinical development of such inhibitors has been hindered by off-target toxicities, poor pharmacodynamics, and lack of target specificity^3,4,6,15^. Thus, identifying novel small-molecule modulators of ERAD is crucial for developing effective therapies for MM, particularly in R/R cases.

To identify small molecules that modulate ERAD for therapeutic and research purposes, we developed a high-throughput cell-based drug screening approach and screened the FDA repurposing library. Here, we report the identification of omaveloxolone (RTA408) as a novel ERAD inhibitor. RTA408 selectively inhibits the degradation of luminal and membrane ERAD substrates but not cytosolic proteins. In MM cells, RTA408 induces UPR activation and rapidly triggers caspase-8-mediated apoptosis via the extrinsic apoptotic pathway, dependent on FADD and RIPK. Originally developed to enhance NRF2 activity by inhibiting its degradation via the KEAP1-CUL3 E3 ligase, RTA408 is FDA-approved for Friedreich’s ataxia^16,17^. However, our findings indicate that RTA408 exerts its cytotoxic effects on MM cells independently of KEAP1, suggesting an ERAD-specific mechanism distinct from NRF2 activation. Importantly, RTA408 is effective against MM cell lines, primary neoplastic plasma cells from MM and plasma cell leukemia patients and *in vivo* MM models independent of PI sensitivity. These findings provide a foundation for preclinical development of ERAD-targeted therapies to improve treatment options for R/R MM.

## RESULTS

### Identification of ERAD Substrate Degradation Inhibitors

To identify ERAD inhibitors, we utilized a GFP-tagged null Hong Kong variant of alpha-1 antitrypsin (NHK-GFP), a well-established ER-retained substrate used to measure ERAD activity^18^. K562 cells were transduced with a lentiviral vector that expresses doxycycline inducible NHK-GFP to develop a stable cell line. Doxycycline was added to the culture for 16h to induce NHK-GFP expression. The cells were then incubated with small molecule compounds in the presence of 20 µM emetine to block transcription and translation, allowing for measurement of NHK-GFP steady state degradation. As positive controls, a VCP/p97 inhibitor, NMS873, and a PI, MG132, were used to block degradation of ERAD substrates. To develop a functional high throughput screen (HTS), we optimized K562 transduction, doxycycline induction, and flow cytometry timepoints. From this optimization, we achieved a HTS Z’ value of >0.6 for screened compound plates between vehicle control and NMS873 (Supp Fig 1A-D). We screened approximately 2,200 compounds from the FDA repurposing library (Selleckchem) and measured relative GFP changes via automated flow cytometry (Fig 1A). HTS software (Mscreen) was used to identify the primary hits, defined as having ≥3 standard deviations (STDEV) from DMSO control^19^. These were further filtered by ≥20% increase in GFP mean florescence intensity (MFI) from the DMSO control (Fig 1A). A total of 121 compounds met these criteria and were validated in triplicate at the same initial testing concentration of 10 µM. Among these, 53 compounds reproducibly inhibited NHK degradation in 3 out of 4 replicates (including the initial run) and were further validated with dose response curves. Ten compounds exhibited an inhibitory concentration 50 (IC_50_) of ≤20 µM (Fig 1B and Supp Fig 2A). Autofluorescent compounds, for example PHA-665752, that shifted the MFI independent of GFP positivity were removed (Supp Fig 2B). Mechanistic prioritization revealed that 3 of the 10 compounds are known modulators of ERAD substrate degradation, including two PIs, bortezomib (BOR) and epoxomicin, and the VCP/p97 inhibitor NMS873 which was used as the positive control in our screen (Supp Fig 2A). Five compounds were previously tested in early phase clinical trials or pre-clinical studies in MM but have not been linked to ERAD activity^20–25^ (Supp Fig 2A). We identified two compounds that inhibited ERAD substrate degradation and had not been characterized in MM, omaveloxone (RTA408) and zinc pyrithione, and selected them for further validation.

**Figure 1:**
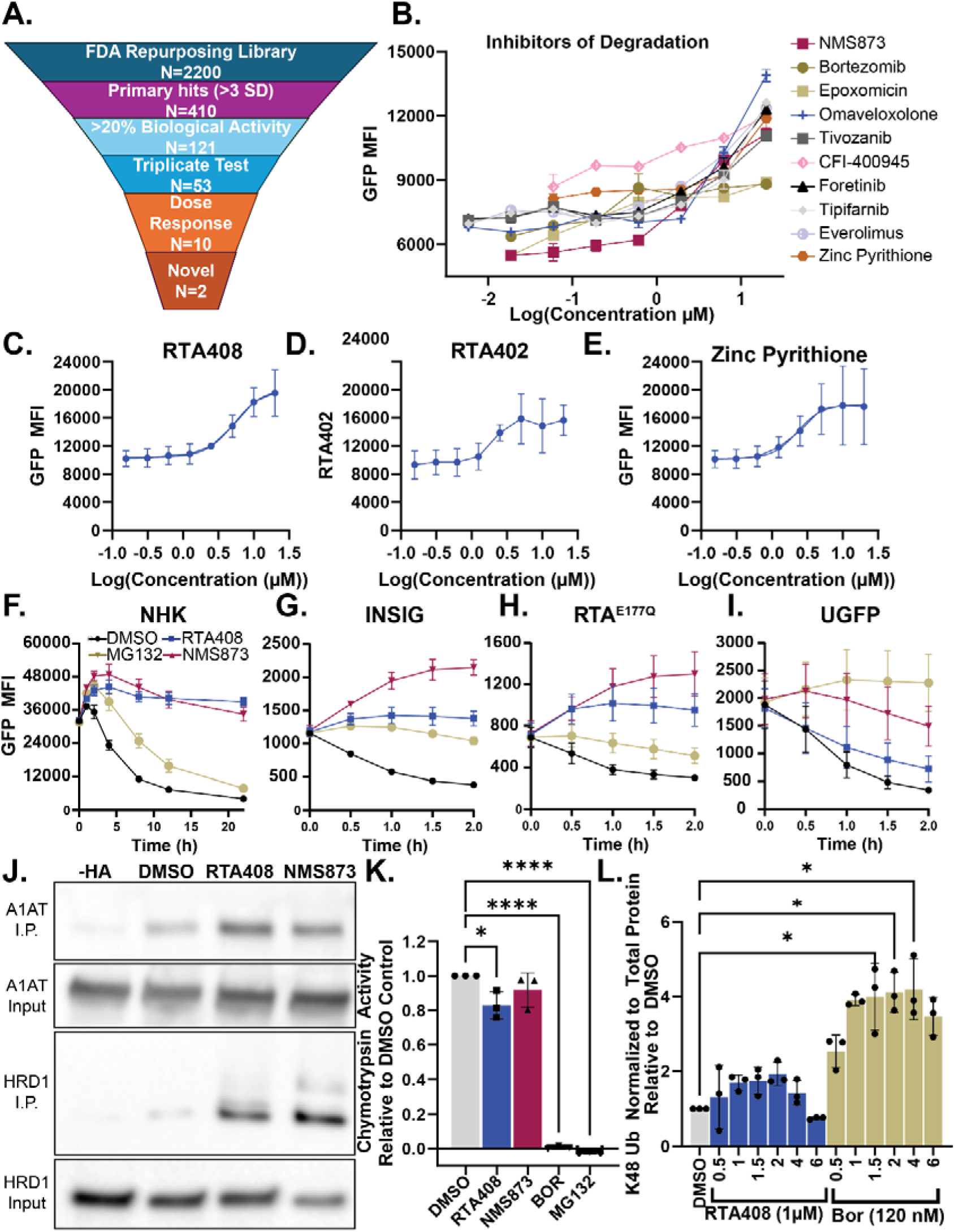
Identification of Inhibitors of ERAD Substrate Degradation. A. NHK-GFP HTS Triage Chart. B. Dose response curve for steady state NHK-GFP degradation in K562 cells treated with small molecule inhibitors at 5.8 nM-20 µM at 4h (N=2 technical replicates). C-E. Dose response curve for inhibition of NHK-GFP steady state degradation by RTA408 (C), RTA402 (D), or zinc pyrithione (E) at 156 nM-20 µM at 4h in K562 cells. F-I. Steady state degradation of NHK-GFP (F), INSIG-GFP (G), RTA^E177Q^-GFP (H), and uGFP (I) with DMSO or 10 µM RTA408, NMS873 or MG132 between 0-2h in K562 cells. J. Immunoprecipitation with HA-UB and detection of alpha-1 antitrypsin (A1AT) and HRD1 by immunoblot following 4h steady state degradation with DMSO or 10 µM RTA408 or NMS873 in K562 cells. K. Chymotrypsin activity measured by cell-based proteasome-Glo™ following 2h treatment with DMSO, 1 µM RTA408, 10 µM NMS873, 120 nM BOR, or 10 µM MG132 in MM.1S. L. Relative quantitation of immunoblotting for whole cell K48 ubiquitination with 1 µM RTA408 or 120 nM BOR treatment from 0.5-6 h in MM.1S cells. All steady state degradation experiments were performed with 20 µM emetine. N=3 unless specified. Mean±STDEV. Statistical analysis performed with ordinary one-way ANOVA with Dunnett’s multiple comparisons test (Fig 1K) and Kruskal-Wallis test (Fig 1L). *p≤0.05, ****p≤0.0001

**Figure 2.**
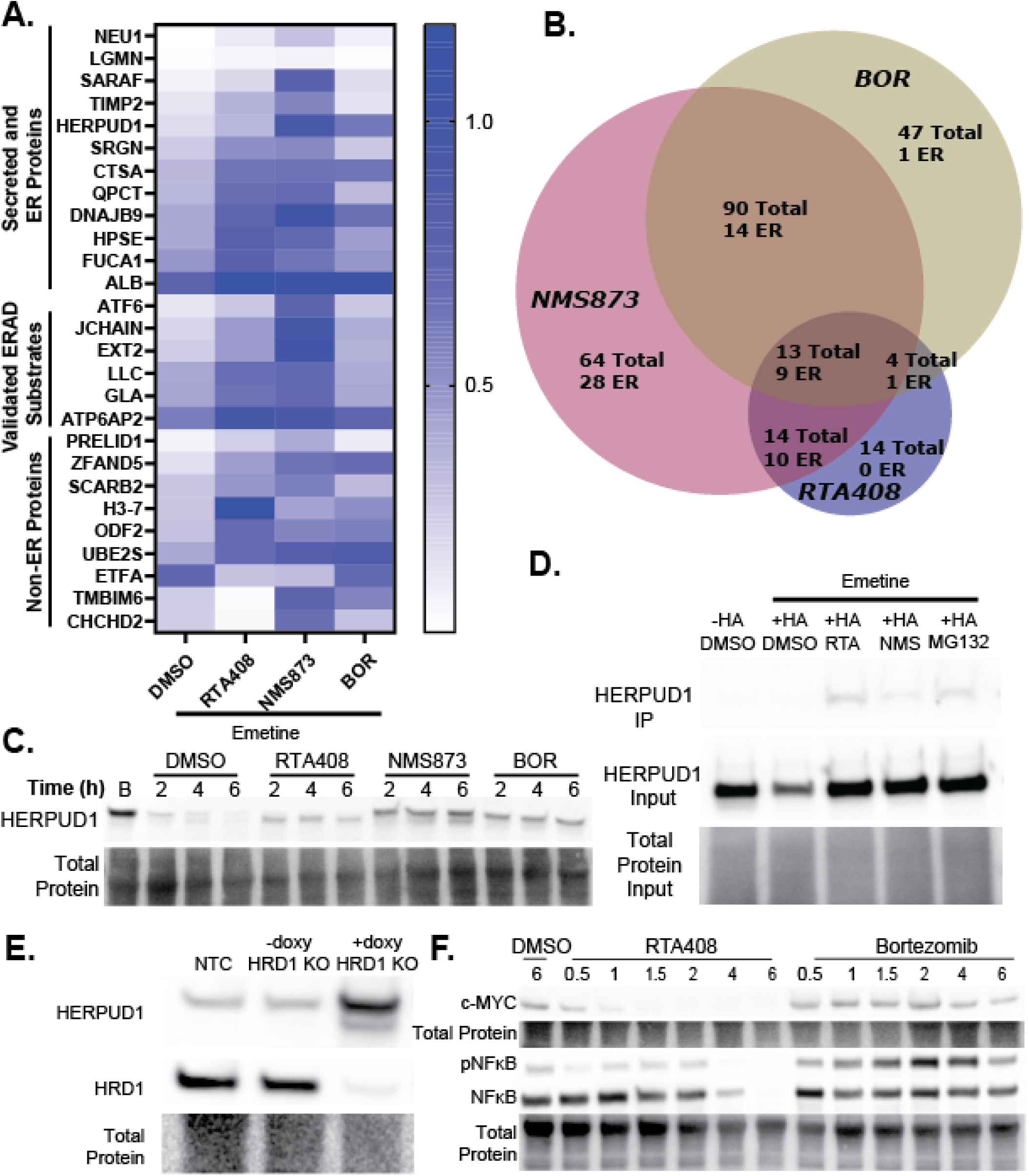
Proteomic Analysis with ERAD Inhibition. A. Heatmap of relative protein abundance ratios normalized to DMSO control (no emetine) for steady state degradation (50 µM emetine) with DMSO, 1 µM RTA408, 10 µM NMS873, or 120 nM BOR in MM.1S at 4h. B. Venn Diagram summarizing the number of proteins that had significantly altered steady state degradation (adjusted p value ≤0.01) compared to DMSO+emetine control. C. HERPUD1 immunoblot steady state degradation (50 µM emetine) with DMSO, 1 µM RTA408, 10 µM NMS873, or 120 nM BOR in MM.1S. D. Immunoblot for HERPUD1 from immunoprecipitant (IP) or input from HA-tag K562 cells transduced with HA-ubiquitin and treated with DMSO, 10 µM RTA408, NMS873, or MG132. E. Immunoblot of MM.1S transduced with non-targeting control (NTC) or doxycycline inducible HRD1 KO ± doxycycline. F. Immunoblot of MM.1S cells treated with 1 µM RTA408 or 120 nM BOR for 0.5-6h. Immunoblots are representative of N=3-4 replicates.

For validation, commercially available compounds were acquired, and dose response curves were repeated for RTA408 (Fig 1C), bardoxolone methyl (RTA402), the parent compound for RTA408 (Fig 1D), and zinc pyrithione (Fig 1E), all of which showed an IC_50_ for NHK-GFP degradation in the low micromolar range in K562 cells. We also included two established VCP/p97 inhibitors (NMS873 and CB5083; Supp Fig 2C-D) as positive controls for ERAD inhibition. Further analysis demonstrated dose-dependent inhibition of additional ERAD substrates, including the membrane protein INSIG and the non-glycosylated luminal substrate RTA^E177Q^ by both RTA408 and RTA402 (Supp Fig 2E-F). There was minimal inhibition of degradation for the cytosolic protein unstable GFP (uGFP) by RTA408 at less than 10 µM (Supp Fig 2G). However, the parent compound RTA402 inhibited uGFP degradation at 5 µM. Zinc pyrithione exhibited limited inhibition of INSIG and non-glycosylated substrate degradation and was not pursued further (Sup Fig 2E-G).

We found that RTA408 extends the half-life of steady state NHK-GFP degradation from 4.8h to >21h as compared to the proteasome inhibitor MG132, which increases the half-life of NHK-GFP to 10h (Fig 1F). Similarly, RTA408 inhibits the steady state degradation of INSIG and RTA^E177Q^ over 2h, whereas there is minimal effect on uGFP degradation as compared to NMS873 or MG132, which prolonged the half-life of uGFP (Fig 1G-I). To assess the effects of RTA408 on ERAD substrate ubiquitination, K562 cells were transduced with HA-tagged ubiquitin and subjected to steady state degradation in the presence of RTA408 or NMS873. We found that NHK-GFP and HRD1 remain ubiquitinated in both conditions (Fig 1J), indicating RTA408 does not interfere with the ubiquitination of ERAD substrates and likely inhibits activity downstream of HRD1. Furthermore, RTA408 has minimal effects on chymotrypsin activity in the proteasome as compared to the BOR or MG132 (Fig 1K), indicating RTA408 does not directly inhibit proteasomal degradation. Similarly, RTA408 treatment does not significantly increase total K48-linked ubiquitination or overall ubiquitination levels, whereas BOR induces a pronounced accumulation of ubiquitinated proteins (Fig 1L and Sup 2H-I). Together, these findings suggest that RTA408 inhibits degradation of both luminal and membrane ERAD substrates but has minimal effect on the degradation of select cytosolic proteins in a manner that is independent and distinct from proteasome inhibition.

### Modulation of ERAD Substrate Degradation

To gain insight into potential functional overlap between RTA408, NMS873, and BOR, we assessed their effects on steady state protein degradation by quantitative proteomics. Given the variation in substrate regulation across different cell types and our focus on MM, we conducted these analyses using the MM.1S cell line. MM.1S cells were treated with RTA408, NMS873, and BOR for 4h in the presence of 50 µM emetine (Fig 2A-B; Supplemental Data S1-6) to identify proteins whose steady state degradation is modulated by these treatments. We observed a significant difference in abundance of 45 proteins with RTA408 treatment compared to DMSO control (Fig 2B). Of the 45 proteins, 44% are annotated in UniProt as ER proteins or contain an ER signal sequence, whereas only 33% and 16% of proteins altered by NMS873 or BOR, respectively, were ER proteins^26^. Consistent with its known functions, most of the targets of BOR are cytosolic proteins. There was a greater overlap between the ER proteins stabilized by RTA408 and NMS873 than those altered by BOR. Among the proteins shared between RTA408 and NMS873 treatments, 70% were ER proteins, including six validated ERAD substrates, such as lambda light chain and J-chain. Immunoblot analysis confirmed the proteomics findings, showing an inhibition of steady-state lambda light chain degradation between 2-6 hours with RTA408 and NMS873 treatment (Sup Fig3A-B) compared DMSO.

In addition to the known ERAD substates, our proteomic findings also identified 12 additional secreted or ER proteins stabilized by both RTA408 and NMS873. We hypothesized that RTA408, along with NMS873, could be utilized to identify novel ERAD substrates. One such substrate is HERPUD1, which was stabilized by treatment with RTA408, NMS873 and BOR according to proteomic analysis (Fig 2A). The stabilization of HERPUD1 was validated by immunoblot (Fig 2C and Supp 3C), albeit the effect of RTA408 did not reach statistical significance. To determine whether HERPUD1 undergoes ubiquitination and degradation via the ubiquitin-proteasome system, we performed HA-immunoprecipitation of K562 cells transduced with HA-ubiquitin. Immunoblot of the immunoprecipitant confirmed HERPUD1 ubiquitination in the presence of RTA408, NMS873, and BOR (Fig 2D). Furthermore, we observed accumulation of HERPUD1 protein levels with genetic KO of HRD1 in MM.1S, which ablates ERAD activity (Fig 2E and Supp Fig3D). Altogether these results suggest that RTA408 alters ER protein and ERAD substrate degradation and may be utilized as a tool to identify novel ERAD substrates.

We also observed differential regulation of cytosolic substrates between RTA408 and BOR in MM cells. We detected a decrease in c-MYC levels, which is implicated in MM proliferation and progression, following RTA408 treatment whereas the levels are stabilized with BOR treatment (Fig2F and Supp Fig3E)^27^. Similarly, BOR has been reported to inhibit IkB degradation, leading to increased phosphorylation and nuclear translocation of NFκB, a transcription factor that plays a key role in the survival and proliferation of MM^28,29^. Following RTA408 treatment, we observed decreased pNFκB whereas BOR leads to a moderate increase in NFκB phosphorylation (Fig2F and Supp Fig3F). These findings highlight the differential effects of RTA408 and BOR on cytosolic protein regulation, underscoring the distinct consequences of alternative ERAD inhibition strategies on both ER and cytosolic protein homeostasis.

### Cytotoxicity of ERAD Inhibition in Multiple Myeloma Cell Lines

Previous studies have suggested that ERAD inhibition via PI or p97/VCP inhibitors contributes to MM cytotoxicity by triggering ER stress^4,9^. We sought to test whether ERAD inhibition with our novel small molecule inhibitors were cytotoxic to MM cells. Treatment with RTA408 decreased cell viability in nine different multiple myeloma cell lines within 24 h of treatment with IC_50_ values ranging from 104-489 nM (Fig 3A and Supp Fig 4A). Cytotoxicity occurred in a time dependent manner within 12h (Fig 3B and Supp Fig 4A) with maximal effects observed within 72 hours (Fig 3C and Supp Fig4A). The decrease in MM cell viability was confirmed with alternative viability measurements using Calcein AM esterase-based staining (Fig 3D) or live/dead nuclear staining by flow cytometry (Supp Fig4B). Similarly, RTA402 is also cytotoxic to MM cell lines at 24h (Fig 3E). To determine whether RTA408-mediated cytotoxicity is dependent on its known target KEAP1, we performed inducible CRISPR-Cas9 knockout (KO) of KEAP1 in MM cells. KEAP1 KO did not alter RTA408 cytotoxicity (Fig3F; Supp Fig 4A, 4C, and Supp Fig 5A-B), suggesting that RTA408-induced MM cell death is independent of KEAP1.

**Figure 3.**
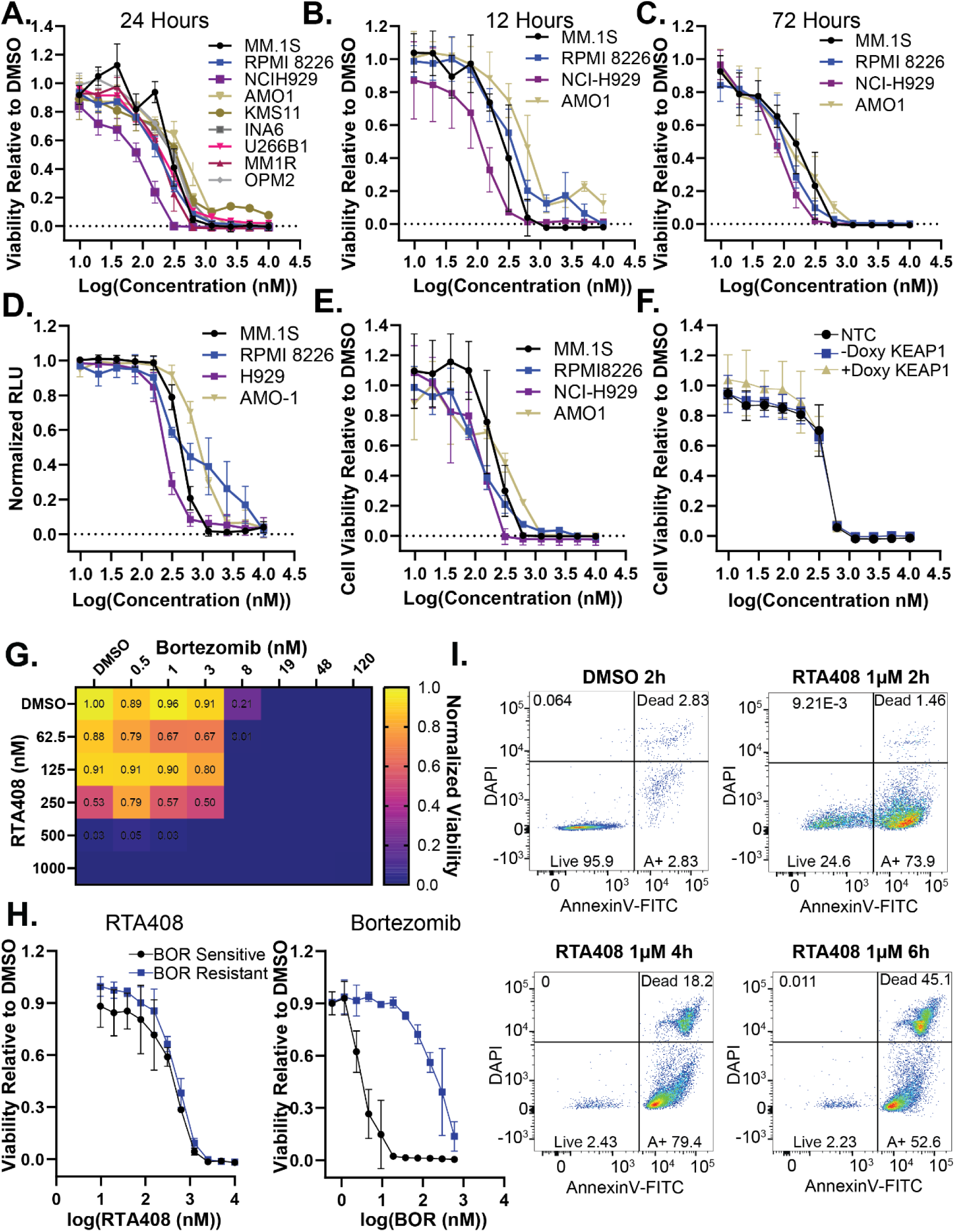
RTA408 cytotoxicity in Multiple Myeloma Cell Lines. A-C. MM cell line viability measured by CellTiter-Glo® following treatment with 10 nM-10 µM RTA408 for 24h (A), 12h (B), and 72h (C). D. Calcein AM uptake in MM cell lines treated with 10 nM-10 µM RTA408 for 24h. E. MM cell line viability measured by CellTiter-Glo® following treatment with 10 nM-10 µM RTA402 for 24h. F. MM cell line viability measured by CellTiter-Glo® in MM.1S cells with non-targeting control (NTC) or a doxycycline inducible KEAP1 KO ±doxycycline induction treated with 10 nM-10 µM RTA408 for 24h. G. Heatmap of mean viability measured by CellTiter-Glo® in MM.1S cells 24h following Bortezomib 0.5-120 nM and RTA408 62.5-1000 nM cotreatment. Viability is normalized to DMSO control. H. Representative flow cytometry plots of AnnexinV-FITC and DAPI staining in MM.1S cells treated with DMSO or 1 µM RTA408 at respective timepoints. N=3. Mean±STDEV.

**Figure 4.**
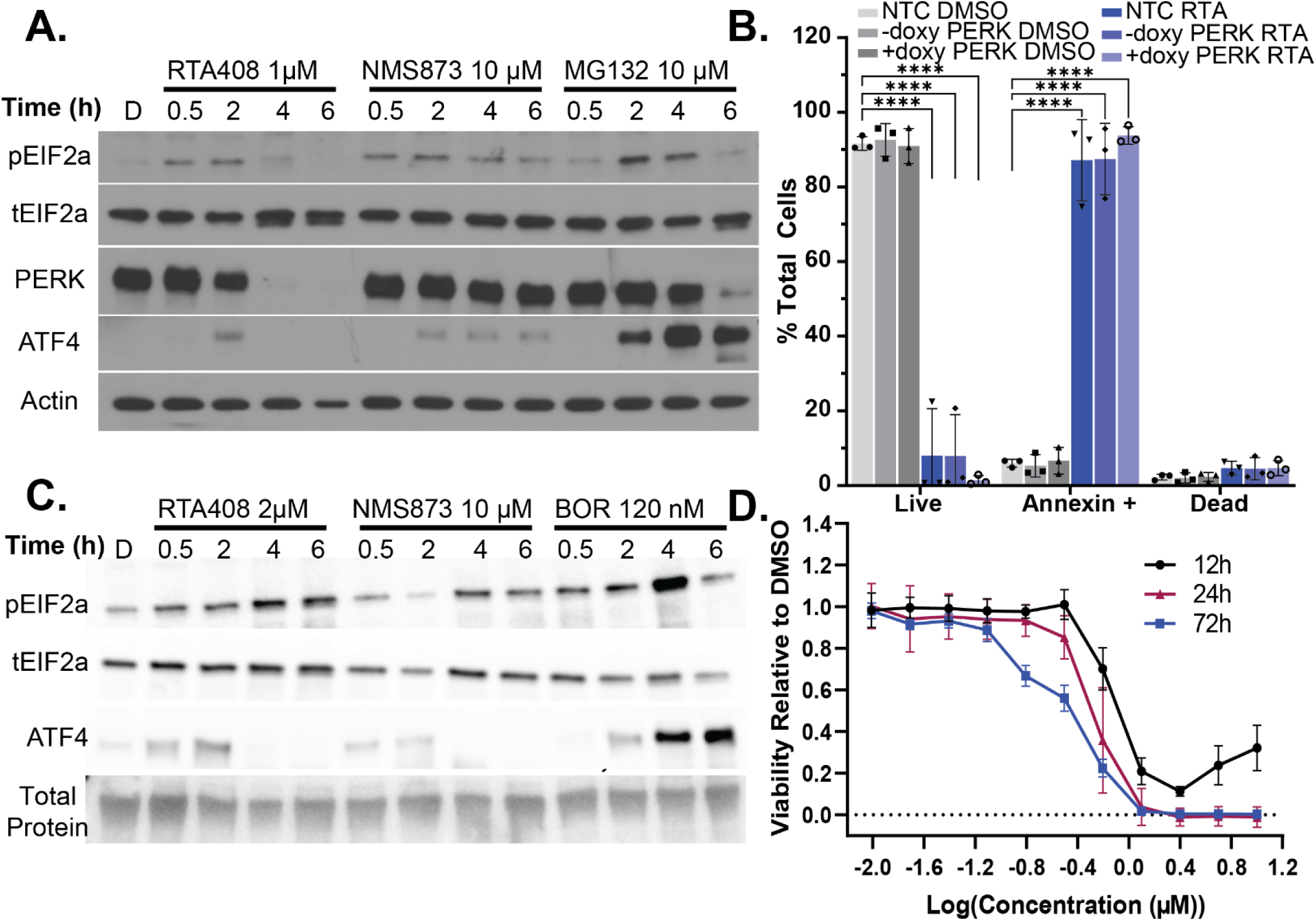
Cell death induced by ERAD inhibition is independent of UPR activation or immunoglobulin hypersecretion. A. Immunoblot of pEIF2a, total EIF2a, PERK, ATF4, and β-Actin in MM.1s treated with 1 µM RTA408, 10 µM NMS873, or 10 µM MG132 for 0.5-6h. B. Quantitation of live (AnnexinV-DAPI-), Annexin+ (AnnexinV+DAPI-) or dead (AnnexinV+DAPI+) population 4h in MM.1S cells transduced with non-targeting control, or doxycycline inducible PERK KO ± doxycycline. C. Immunoblot of pEIF2a, total EIF2a, ATF4, and total protein quantitation in KMS12BM treated with 2 µM RTA408, 10 µM NMS873, or 120 nM BOR for 0.5-6h. D. KMS12BM cell line viability measured by CellTiter-Glo® following treatment with 10 nM-10 µM RTA408 for 12-72h. Immunoblots are representative of N=3-4 replicates. Mean±STDEV. Statistical analysis performed with a two-way ANOVA with Tukey’s multiple comparison test. **** p<0.0001

**Figure 5.**
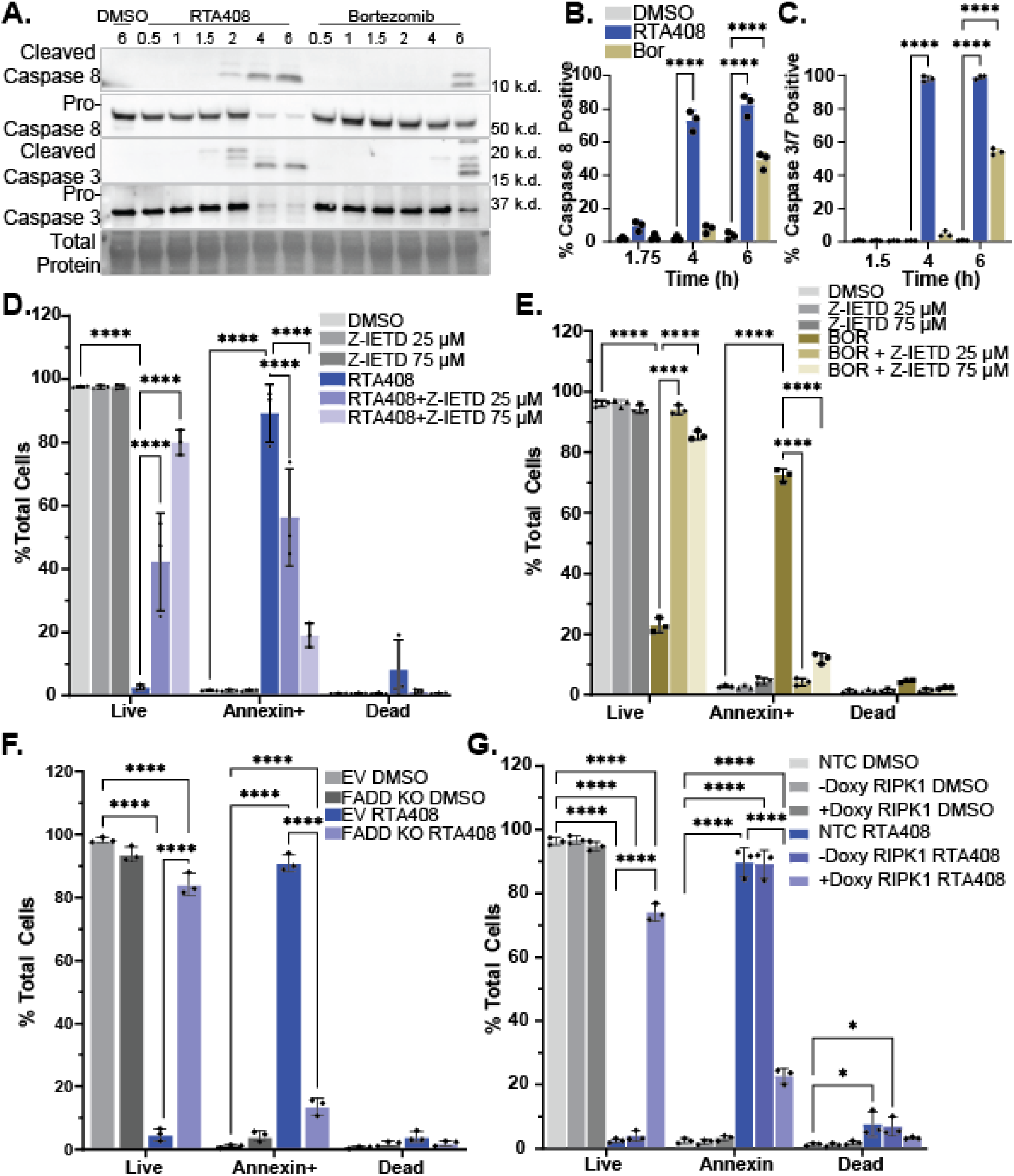
Pro-Apoptotic Signaling with ERAD Inhibition. A. Immunoblot of caspase 8 and 3 (cleaved and pro-forms) and total protein in MM.1s treated with DMSO, 1 µM RTA408 or 120 nM BOR for 0.5-6h. B-C. Flow cytometry quantitation of caspase 8 (B) or caspase 3/7 (C) activity in live MM.1S following DMSO, 1 µM RTA408 or 120 nM BOR for 1.5-6h. D-E. Flow cytometry quantitation of live (AnnexinV-DAPI-), Annexin+ (AnnexinV+DAPI-) or dead (AnnexinV+DAPI+) in MM.1S treated with 25-75 µM Z-IETD-FMK and DMSO or 1 µM RTA408 for 4h (D) or 120 nM BOR for 6h (E). F-G. Flow cytometry of Annexin V staining in MM.1S cells transduced with empty vector (EV), non-targeting control (NTC), or doxycycline inducible FADD KO (F) or RIPK1 KO (G) treated with DMSO or 1 µM RTA408 for 4h. N=3. Mean±STDEV. Statistical analysis performed with a two-way ANOVA with Tukey’s multiple comparison test. *P<0.05 ****p≤0.0001

Next, we evaluated the potential combinatorial effects of RTA408 with other MM therapies. Co-treatment with BOR did not enhance RTA408-induced cytotoxicity in MM.1S cells (Fig 3G). RTA408 cytotoxicity did not correlate with BOR sensitivity across different cell lines (Supp Fig 4B) and was not altered in a BOR-resistant AMO1 cell line (Fig 3H and Supp Fig 4A). In contrast, co-treatment with lenalidomide (Supp Fig 4D) or dexamethasone (Supp Fig 4E) exhibited a combinatorial effect. These data suggest that RTA408 cytotoxicity is independent of PI sensitivity.

Given the similar IC_50_ between 12 and 24h treatment, we sought to determine the timing of cytotoxicity in MM cells. We observed that RTA408 led to an early induction of apoptosis (1.5-2 h) measured by Annexin V positivity followed by an increase in DAPI uptake within 4-6h of treatment (Fig 3I and Supp Fig 4F-H). In contrast, the induction of Annexin V positivity and DAPI uptake did not occur until after 6h with NMS873 or BOR. These data suggest that RTA408 induces a rapid induction of apoptosis and cell death in MM cells.

### Induction of the Unfolded Protein Response by ERAD Inhibition in MM Is Independent of Apoptosis

Inhibition of ERAD activity has previously been associated with the rapid induction of ER stress and activation of the UPR. Indeed, treatment of MM.1S cells with RTA408, NMS873 or MG132 led to a rapid induction of pEIF2α and accumulation of ATF4 protein (Fig 4A, Supp Fig 6A). Similar activation of PERK signaling was observed with RTA402, CB5083, and BOR treatment (Supp Fig 6B) as well as in RPMI8226 cells (Supp Fig 7A-B). Increased mRNA expression of CHOP was detected within 2-4h of treatment (Supp Fig 6C and Supp Fig 7C), although we were unable to detect CHOP protein (data not shown). There is also an increase in IRE1α signaling with an increased ratio of spliced/total XBP1 at 2 h in MM.1S (Supp Fig6D) and RPMI8226 (Supp Fig 7D) cells. IRE1α has been reported as an ERAD substrate in genetic models with ERAD deficiency, however, we did not observe a significant increase of IRE1α levels in MM.1s (Supp Fig 6E) or RPMI8226 (Supp Fig 7E) cells with chemical inhibition of ERAD.

**Figure 6.**
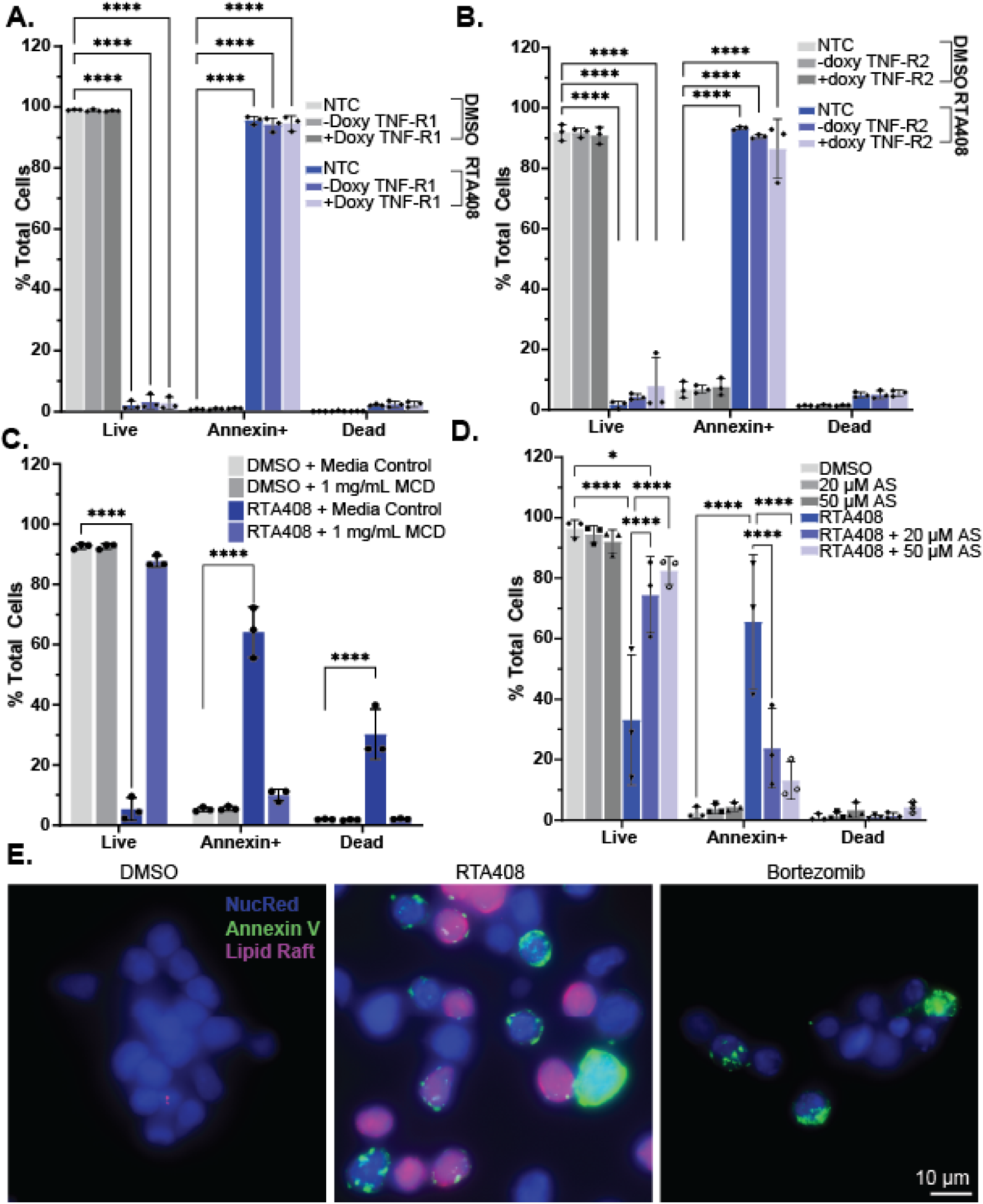
Lipid Raft Dependent Activation of Apoptosis with ERAD Inhibition. A.-B. Flow cytometry quantitation of live (AnnexinV-DAPI-), Annexin+ (AnnexinV+DAPI-) or dead (AnnexinV+DAPI+) in MM.1S cells transduced with non-targeting control, or doxycycline inducible TNF-R1 (A), TNF-R2 (B) treated with DMSO or 1 µM RTA408 for 4h. C. Flow cytometry quantitation of Annexin V staining in MM.1S treated with 1 mg/mL MCD and DMSO or 1 µM RTA408 for 4h. D. Flow cytometry quantitation of Annexin V staining in MM.1S cells treated with DMSO or 1 µM RTA408 4h and DMSO or 20-50 µM atorvastatin (AS) for 22h. E. Representative immunofluorescence images for MM.1S cells treated with DMSO, 1 µM RTA408, or 120 nM BOR for 2h and stained for NucRed-Alexa647, Vybrant-Lipid Raft Label-Alexa555 (Lipid Raft), or AnnexinV-Alexa488. N=3. Mean±STDEV. Statistical analysis performed with a two-way ANOVA with Tukey’s multiple comparison test. ****p<0.0001

**Figure 7.**
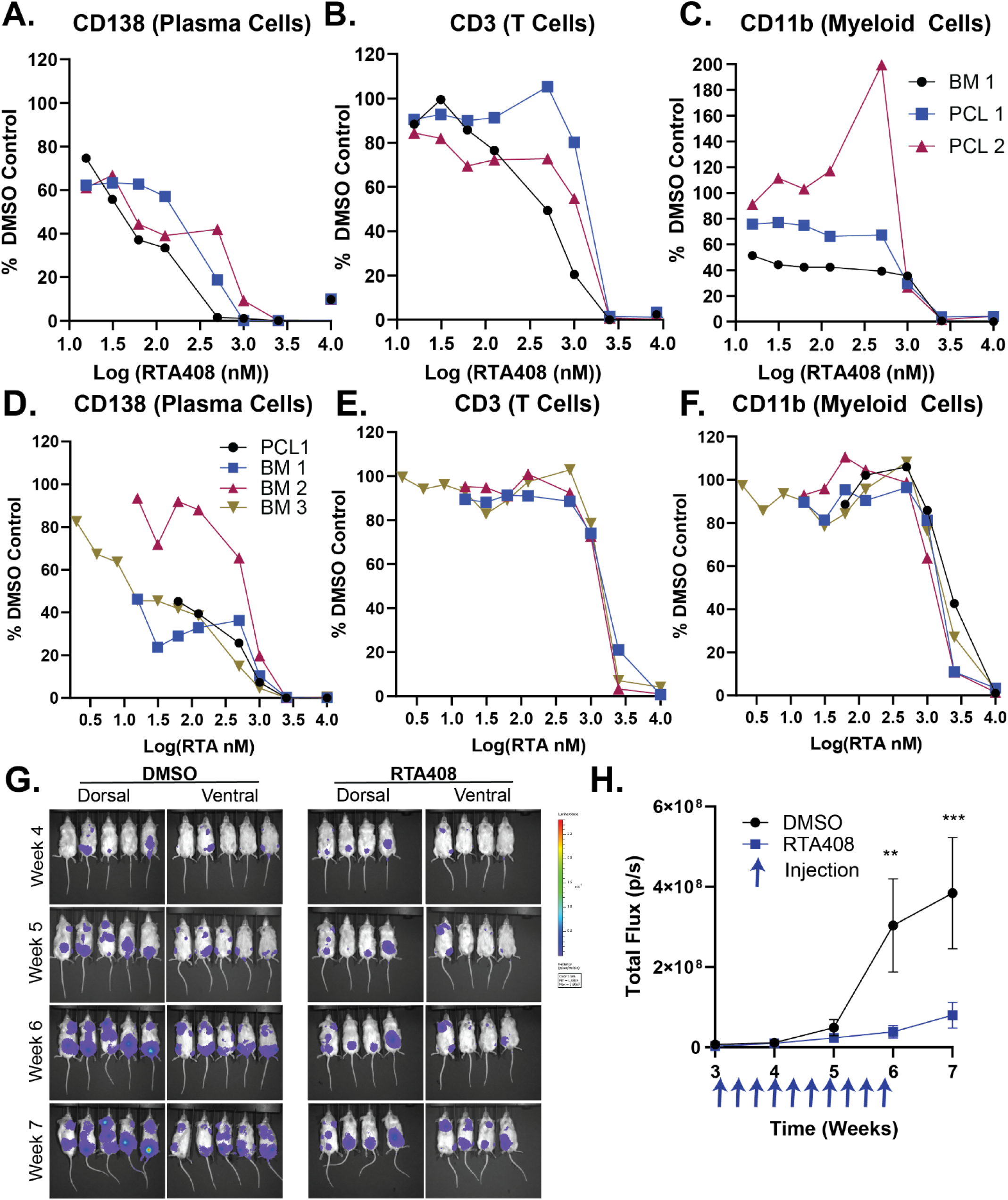
RTA408 Cytotoxicity in Primary Cells and In Vivo: A-C. Flow cytometry analysis of Annexin-DAPI-CD138 (A), CD3 (B), or CD11b (C) cells following 60h RTA408 in primary bone marrow (BM) or peripheral blood (PCL) cells from patients with relapsed refractory MM. D-F. Flow cytometry analysis of Annexin-DAPI-CD138 (D), CD3 (E), or CD11b (F) cells following 36h RTA408 in primary bone marrow (BM) or peripheral blood (PCL) cells from patients with newly diagnosed MM. G-H. Bioluminescent imaging and quantitation of total luciferin flux in NSG mice transplanted with 5e6 RPMI8226-luciferase cells. N=5-6 for RPMI transplant. Mean±STDEV. Statistical analysis performed with a two-way ANOVA with Tukey’s multiple comparison test. ** p<0.01,

While induction of apoptosis has been associated with upregulation of CHOP expression from UPR signaling, it has not been established that the UPR is directly responsible for early apoptotic signaling in response to ERAD inhibition in MM. To test this, we inhibited PERK with either doxycycline inducible CRISPR-CAS9 KO or small molecule inhibitors of PERK signaling (GSK2606414; G414) or pEIF2α activation (ISRIB). Despite moderately decreasing pEIF2α and ATF4 levels (Supp Fig 8A-B, D, F), inhibition of PERK or pEIF2a signaling did not induce a meaningful decrease in annexin positive populations (Fig4B; Supp 8C, E). However, an important limitation to these studies is that there was not complete ablation of pEIF2α or ATF4 accumulation with PERK KO or pharmacologic inhibition, which may be due to activation of the PERK-independent integrated stress response (ISR) or alternative arms of UPR. Next, we utilized treatment with emetine, which inhibits transcription and protein translation, to evaluate if the induction of early apoptosis was dependent on the downstream transcription/translation of stress response proteins. We found that while completely ablating ATF4 accumulation, emetine failed to rescue MM.1S cells from early apoptosis after RTA408 treatment (Supp Fig 8G-I). These studies suggest that the early induction of apoptosis and cytotoxicity from RTA408 is likely independent from transcriptional and downstream stress responses in the UPR in MM.

It has been assumed that the activation of the UPR and ER stress responses are due to the secretory nature of MM cells and high burden of paraprotein production. To further evaluate the contribution of the UPR to RTA408-induced apoptosis, we evaluated KMS12BM cells, which lack paraprotein production at both the mRNA and protein level^30^. We observed that RTA408 rapidly induced activation of pEIF2α and ATF4 accumulation and a time-dependent increase of apoptosis in KMS12BM cells (Fig 4C-D), like secretory MM cell lines. These results suggest that RTA408-induced early apoptosis and cytotoxicity are independent of both UPR signaling and paraprotein production in MM cells.

### ERAD Inhibition Activates Caspase 8 Induced Apoptosis in MM

Given that RTA408-mediated early apoptotic signaling was independent of transcriptional or translational stress responses, we next evaluated the early pro-apoptotic signaling following treatment with ERAD inhibition. BOR has previously been associated with an induction of apoptosis through modulation of both intrinsic and extrinsic pro-apoptotic pathways^31–35^. We observed a rapid induction of cleaved caspase 8 and caspase 3 within 2h of treatment of RTA408 (Fig 5A) and RTA402 (Supp Fig 9A), which precedes activation at 6h in BOR treated MM.1s cells (Fig 5A). This correlates with caspase 8 (Fig 5B) and caspase 3/7 activity (Fig 5C and Supp Fig 9B) in MM1S. Activation of caspase 8 and caspase 3/7 was also observed with RTA408 in RPMI8226 cells (Supp Fig 10A-B) and with HRD1 KO in MM.1S (Supp Fig 10C). We found that the induction of early apoptosis with Annexin V positivity by RTA408 at 4h was blocked by the addition of a pan-caspase inhibitor, Z-VAD-FMK, (Supp Fig 9C) or a caspase 8 specific inhibitor, Z-IETD-FMK, in both MM.1S (Fig 5D) and RPMI8226 cells (Supp Fig 10D). We observed similar reversal of Annexin V positivity at 6h with BOR treatment (Fig5E and Supp Fig 9D). These results indicate that early apoptosis from the loss of ERAD activity is mediated by caspase 8 activation.

Caspase 8 activation and pro-apoptotic signaling have been classically associated with formation of the death-inducing signaling complex (DISC) and activation of the extrinsic apoptotic pathway (Supp Fig 12H)^31,36^. To determine whether DISC components are required for RTA408-induced apoptosis, we performed CRISPR-Cas9 knockout of FADD, an essential adapter protein in caspase-8 mediated apoptosis. FADD KO rescued early apoptosis induction at 4h with RTA408 treatment (Fig 5F; Supp 5C) and provided a partial rescue with a 20% increase of live cells at 6h with BOR treatment (Supp Fig 9E). Similarly, we observed significant rescue from early apoptosis with inducible knockout of RIPK1 after 4h of treatment with RTA408 (Fig 5G; Supp 5D) and 6h of treatment with BOR (Supp Fig 9F). These data implicate RIPK1 and FADD mediated activation of caspase 8 and downstream pro-apoptotic signaling as important mediators of RTA408-induced apoptosis in MM cells.

### Cholesterol Dependent Caspase 8 Activation with ERAD Inhibition

Signaling through various cell death receptors has been associated with DISC signaling through the activation of FADD and RIPK1, which can ultimately lead to the induction of apoptosis through caspase 8 (Supp Fig 12H)^37–39^. To further characterize the mechanism of RIPK1-FADD activation following RTA408 treatment, we investigated the role of FAS and tumor necrosis factor receptors (TNF-R1 and TNF-R2), which have been implicated in apoptosis induction^40^. While we detected cleaved RIPK1 within 1.5h of treatment with RTA408 and 4h with BOR; FAS, TNF-R1, or TNF-R2 levels remained did not increase (Supp 11A). Additionally, inducible CRISPR-CAS9 KO of FAS, TNF-R1, or TNF-R2 (Supp Fig 5E-F) failed to rescue the early apoptotic signaling in MM.1S treated with RTA408 (Fig 6A-B; Supp Fig 11B) or BOR (Supp Fig 11C-E). Single knock out of additional extrinsic cell death receptors DR4 and DR5 similarly failed to rescue early apoptotic signaling induced by RTA408 or BOR (Supp Fig 5G-H and 11F-G)^41^ nor did we observe an accumulation of FADD following treatment (Supp Fig11A).

Given that we were unable to identify an individual cell death receptor responsible for pro-apoptotic signaling, we evaluated alternative mechanisms for intracellular activation of RIPK1, FADD, and caspase 8 signaling^42–46^. Treatment with cathepsin inhibitors Gly-Phe-β-Naphthylamide (GPN) or Pepstatin A did not rescue RTA408 mediated apoptosis (Supp Fig 12A-B). We also observed that the early induction of apoptosis was independent of KEAP1 (Supp Fig 12C). Because IRE1α has been previously shown to activate RIPK1 signaling directly^47^, we performed CRISPR-CAS9 knockout of IRE1α^46^. IRE1α KO failed to restore the live population following the with RTA408 (Supp 5I, Supp12D) or BOR (Supp Fig 12E). Finally, while activation of RIPK1 has been associated with reactive oxygen species (ROS) production, ERAD inhibition did not result in a significant increase in ROS levels (Supp 12F-G).

To evaluate whether apoptotic induction is reliant on membrane associated signaling and lipid raft organization, we treated cells with methyl-β-cyclodextrin (MCD), a compound known to deplete cholesterol from the plasma membrane, which disrupts both lipid rafts and DISC assembly^48^. Co-treatment with MCD restored the live cell population after 4h RTA408 treatment in MM.1S (Fig 6C) and RPMI8226 (Supp Fig 10E) but failed to restore the live cell population following 6h treatment with BOR in MM.1S (Supp Fig 13A) or RPMI8226 (Supp Fig 10F). Similarly, MCD treatment prevented caspase 8 and caspase 3 cleavage and partially restored cell viability at 12h in RTA408- and NMS873-treated cells (Supp Fig 13B-C) but not in BOR-treated cells. Notably, MCD treatment did not affect NHK-GFP degradation in K562 cells and only partially rescued pEIF2α levels in MM.1S (Supp Fig 13 D-F), indicating that RTA408 retained its effect on ERAD activity in the presence of MCD, but it’s effects on pro-apoptotic signaling were ablated. This implies that while RTA408 and PIs induce pro-apoptotic caspase 8 signaling though distinct mechanisms and that the effects of RTA408 are dependent on membrane associated DISC signaling.

Given that ERAD has been implicated in regulation of cholesterol synthesis, lipid raft proteins, and triglyceride homeostasis, we next investigated whether RTA408 altered intrinsic activation of the DISC or lipid raft organization. We found that inhibition of endogenous cholesterol synthesis with atorvastatin (AS) prevented the early induction of apoptosis at 2h and 4h with RTA408 and at 6h after BOR 6h treatment, without affecting the RTA408-induced pEIF2α activation (Fig 6D; Supp Fig13 G-J). Similarly, we observed increased staining and confluence of the lipid raft associated ganglioside GM1 by cholera toxin B staining in MM.1S treated with RTA408 for 2h as compared to the DMSO control or 6h treatment with BOR (Fig 6E). These data suggest that ERAD inhibition by RTA408 leads to altered lipid raft organization and leads to aberrant DISC activation at the plasma membrane, resulting in the induction of pro-apoptotic signaling in MM cells.

### Cytotoxicity in Primary Samples and *In Vivo MM Models*

We next tested the cytotoxic effects of ERAD inhibition by RTA408 in primary patient samples and *in vivo* MM models. We tested peripheral blood or bone marrow mononuclear cells collected from both patients with relapsed and refractory (R/R) disease at the time of disease progression on PI-containing therapy, and patients with treatment naïve disease. We observed maximal cytotoxicity of CD138^+^ cells within 60h of treatment in R/R samples (Fig 7A; Supp 15A-B). Furthermore, there is differential cytotoxicity when compared to non-malignant CD3+ T-cells (Fig7B; Supp 15A-B) and CD11+ myeloid cells (Fig 7C; Supp 15B) from the same samples. However, intermediate toxicity was observed in the CD19⁺ B-cell population (Supp Fig 15D).

To assess whether there is a difference in susceptibility between newly diagnosed and PI-resistant R/R MM, we tested RTA408 cytotoxicity in primary samples from newly diagnosed patients. While we observed similar dose response in CD138^+^ plasma cells, maximal toxicity occurred earlier within 36h of treatment (Fig 7D; Supp 15C). The non-malignant CD3^+^ (Fig7E; Supp 15C), CD11^+^ (Fig7F; Supp 15C), and CD19^+^ (Supp Fig 15C, E) cells were less sensitive to killing by RTA408, suggesting a potential therapeutic index.

To evaluate the efficacy of RT408 *in vivo*, we utilized a bioluminescent xenograft model with intravenous injection of RPMI8226 cells expressing luciferase into sublethal irradiated NSG mice as previously described^49,50^. Three weeks after injection, baseline imaging was performed, followed by intraperitoneal administration of RTA408 at 5 mg/kg every other day for three weeks. RTA408 treatment effectively prevented tumor growth with three weeks of treatment (Fig 7G-H; Supp 15F-G) as compared to the DMSO control. These data suggest that RTA408 is cytotoxic to primary malignant plasma cells and has *in vivo* anti-myeloma activity.

## Discussion

ERAD is crucial for maintaining ER protein homeostasis by facilitating the translocation and degradation of misfolded or target proteins. Despite its essential role in cellular function and survival, the regulatory mechanisms governing ERAD remain poorly understood^51^. A major challenge in studying ERAD is the limited availability of specific inhibitors that directly target ERAD activity. Current small-molecule modulators predominantly act on downstream pathways, such as VCP/p97 or the proteasome, which are not specific to ER protein degradation^2,7^. In this study, we conducted the first cell-based, high-throughput screen to identify small molecules that modulate ERAD activity. A pilot screen using the FDA-repurposing library successfully identified proteasome and VCP/p97 inhibitors as inhibitors of ERAD substrate degradation, validating the robustness of our screening platform. Our screen also identified RTA408 and its parent compound, RTA402, as novel ERAD inhibitors with distinct properties from VCP/p97 or proteasome inhibitors. This study establishes a valuable framework for identifying additional small-molecule modulators that regulate different ERAD components for research and potential therapeutic applications.

The top hit from our screen, RTA408, is an FDA-approved drug for Friedreich’s ataxia, where its proposed mechanism of action involves inhibiting KEAP1-mediated degradation of NRF2 via the CUL3 E3 ligase, thereby stabilizing NRF2 to induce an antioxidant response^52^. However, our data indicate that Keap1 knockout does not affect the cytotoxicity and pro-apoptotic signaling of RTA408 in MM cells. These findings suggest that RTA408 inhibits ERAD independent of KEAP1-NRF2 regulation. While the specific molecular target and precise mechanism of ERAD inhibition by RTA408 remain to be elucidated, our studies suggest that RTA408 prevents the degradation of luminal, membrane, and non-glycosylated ERAD substrates, while sparing many cytosolic proteins. Our proteomic analysis further highlights the specificity of RTA408 for ER protein substrates, whereas PIs and VCP/p97 inhibitors target a broader range of proteins, including cytosolic substrates.

Despite ERAD’s critical function, its protein substrates are not well characterized and are likely cell- and context-dependent. The small-molecule modulators identified in this study will serve as valuable tools for identifying ERAD substrates. Our global quantitative proteomics analysis revealed that RTA408 inhibits the degradation of six validated endogenous ERAD substrates in MM cells, reinforcing its role in ERAD inhibition. Compared to PIs, RTA408 has greater specificity for inhibiting the degradation of ER proteins (45 vs 11%). Among the 13 ER or secreted proteins whose degradation was inhibited by both RTA408 and NMS873, we validated HERPUD1 as a novel ERAD substrate. HERPUD1 is an ERAD complex component proposed to facilitate the delivery of ubiquitinated ERAD substrates to the proteasome^53,54^. While previous studies have shown HERPUD1 accumulation following knockdown of other ERAD proteins, it had not been definitively classified as an ERAD substrate^54^. Here we provide support that HERPUD1 is ubiquitinated and its degradation is prevented with ERAD inhibitors, confirming that HERPUD1 is an endogenous ERAD substrate. These findings illustrate how RTA408 can be utilized to study ERAD biology and can be used to identify ERAD substrates.

Several ERAD complex proteins have been identified as essential for MM cell survival^12^. Inhibition of ERAD has been proposed as a therapeutic strategy for MM and may contribute to the cytotoxic effects of PIs by inducing ER stress and activating the unfolded protein response (UPR)^9^. However, MM remains an incurable disease, and most patients will develop PI resistance resulting in R/R MM^55^. PI resistance has been associated with altered protein homeostasis, prompting interest in targeting other ERAD components as a therapeutic strategy^55^. Here, we identified and characterized RTA408 as a novel ERAD inhibitor with potent cytotoxic activity against MM cells, independent of PI sensitivity. RTA408 induces rapid cell death within 1.5–2h via caspase-8-mediated pro-apoptotic signaling, highlighting the vulnerability of MM cells to ERAD inhibition. Furthermore, RTA408 demonstrated cytotoxicity in primary malignant plasma cells from both newly diagnosed and R/R MM patients on PI therapy. Importantly, non-malignant T-cells and myeloid cells from the same patients were less affected at these concentrations, suggesting a potential therapeutic window. *In vivo*, RTA408 treatment inhibited tumor growth in a xenograft MM model, supporting its in vivo anti-myeloma activity. Given that RTA408 has been previously tested in human clinical trials, it may be rapidly translated for evaluation in MM patients.

Our findings also emphasize that targeting different ERAD components results in distinct effects on protein homeostasis and cytotoxicity in MM cells. PIs broadly inhibit the ubiquitin-proteasome system, affecting both ERAD substrates and cytosolic proteins. In contrast, RTA408 selectively inhibits ERAD while sparing cytosolic proteins such as c-MYC, which may have implications for disease progression and relapse^56,57^. Additionally, we found that PI-induced caspase 8 signaling is RIPK1-dependent, whereas RTA408-induced apoptosis requires both FADD and RIPK1. RTA408 also alters lipid raft distribution, and its effects can be reversed by methyl-β-cyclodextrin (MCD) treatment, suggesting that while both RTA408 and PIs activate caspase 8-dependent pro-apoptotic signaling, they do so through distinct mechanisms. These differences may explain why RTA408 does not exhibit additive cytotoxicity when combined with BOR yet remains effective against PI-resistant MM models. Future studies should further investigate ERAD substrate regulation and the role of ERAD in caspase-8-dependent pro-apoptotic signaling in PI-resistant MM.

The induction of the UPR, particularly through PERK/ATF4 signaling, has been linked to CHOP transcription and pro-apoptotic signaling^58^. Previous studies have shown that ERAD inhibition, via proteasome and VCP inhibitors, rapidly activates the UPR and triggers ER stress-induced apoptosis ^3,4,9,59^. Our findings indicate that while these pathways are activated in MM cells, the early apoptotic response precedes CHOP activation and occurs independently of PERK signaling or transcriptional and translational stress responses. Instead, RTA408 rapidly induces caspase 8-mediated apoptosis through the death-inducing signaling complex (DISC). Moreover, ERAD-induced cytotoxicity is not dependent on the high secretory burden of MM cells. A non-secretory MM cell line KMS12BM exhibited a similar cytotoxic response to ERAD inhibition by RTA408, challenging the prevailing assumption that MM’s dependence on ERAD is primarily due to its highly active protein synthesis and secretion. This suggests that ERAD inhibition is not cytotoxic to MM cells merely by increasing ER stress from excessive misfolded protein accumulation in the ER.

Interestingly, ERAD inhibition induced early apoptosis was independent of individual death receptors, IRE1α, cathepsin activity, or ROS production, despite their known roles in caspase 8 activation^37–40,42,47,60,61^. Instead, we found RTA408 induced pro-apoptotic signaling is associated with altered lipid raft regulation and can be abrogated by disrupting membrane lipid rafts. This raises the possibility that ERAD inhibition by RTA408 disrupts lipid organization, alters the membrane integration of death receptors, or simultaneously activates multiple death receptors^62–66^. ERAD has been implicated in cholesterol synthesis regulation, triacylglycerol homeostasis, and lipid raft-associated proteins, suggesting that ERAD inhibition may affect lipid metabolism, leading to the intrinsic activation of DISC in MM cells^62,63,67,68^. Future studies should focus on elucidating the precise role of ERAD in lipid raft formation and DISC regulation in MM. Our findings highlight that the ERAD inhibitors identified from our high-throughput screen hold promise not only as potential therapeutics but also as tools to dissect cell type-specific ERAD regulation and downstream signaling.

## Supplemental Material

### Methods

#### Key Resource Table

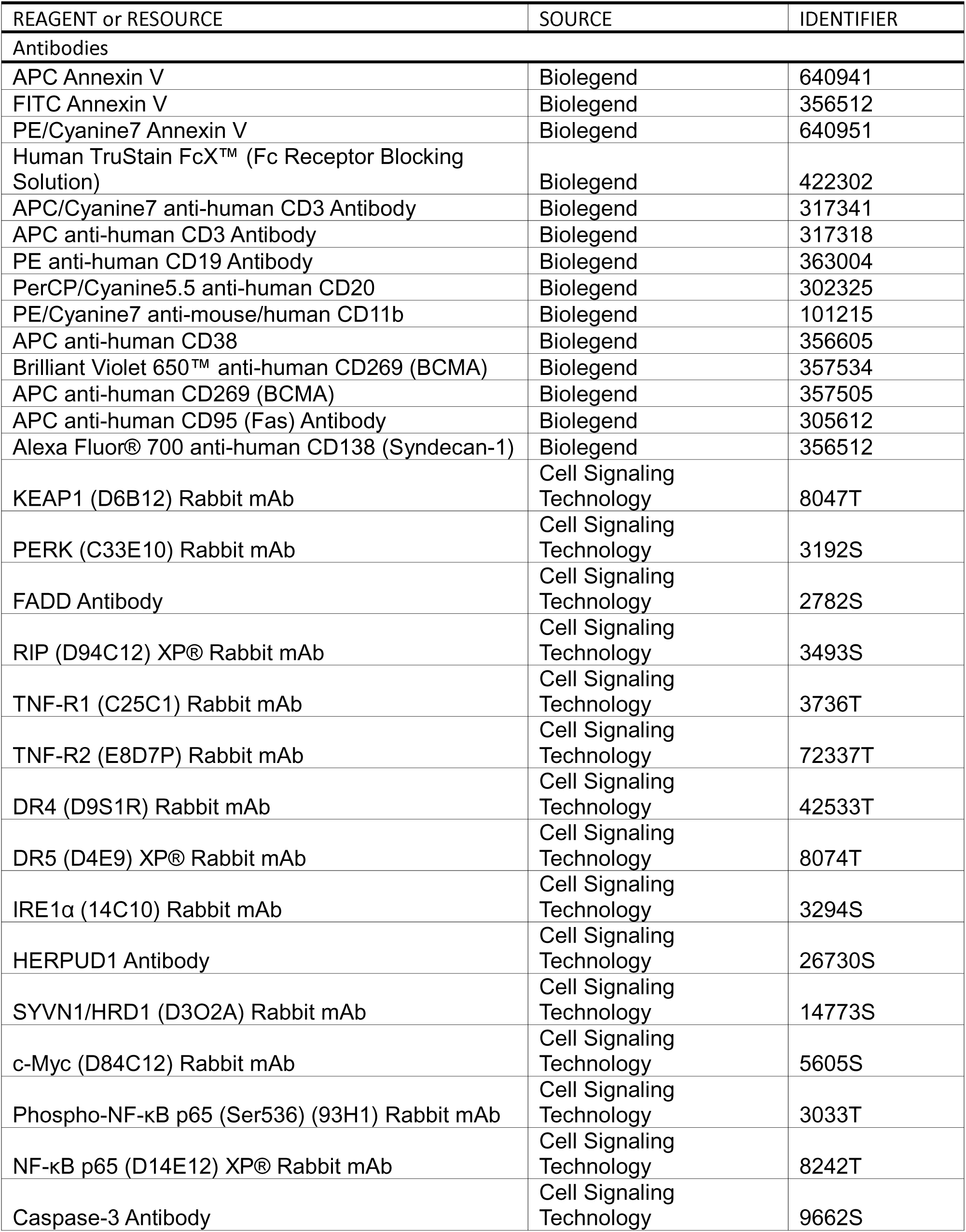
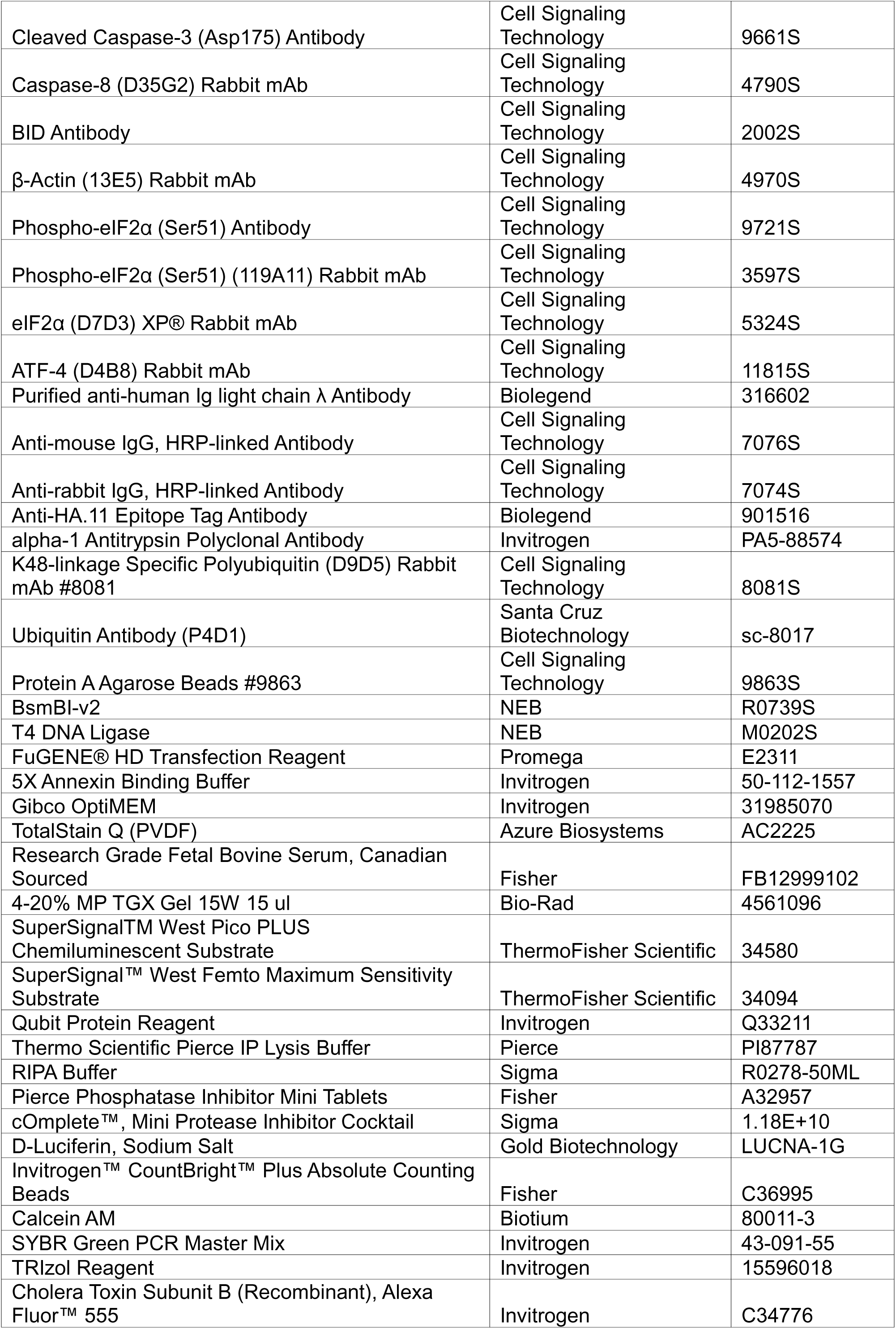
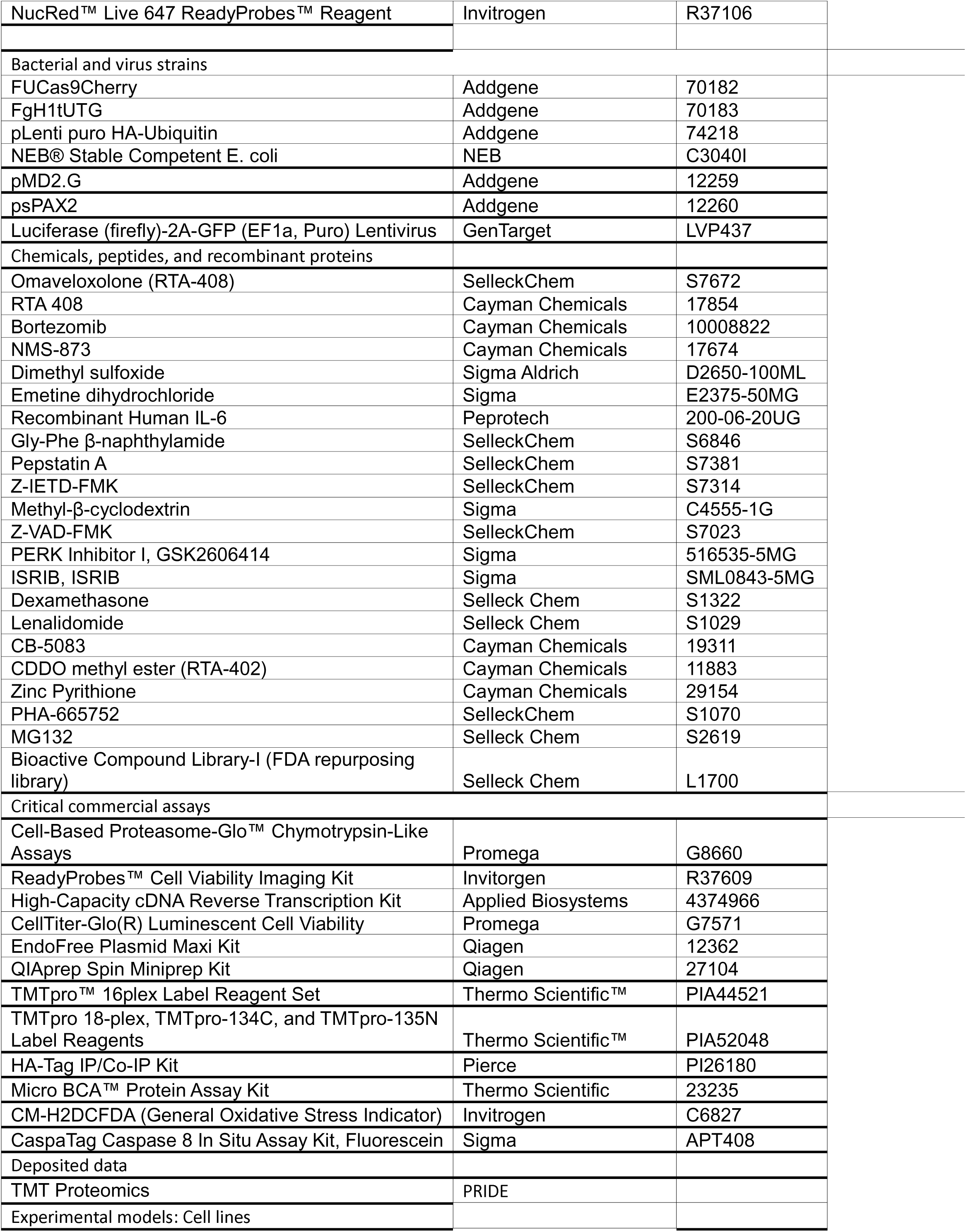
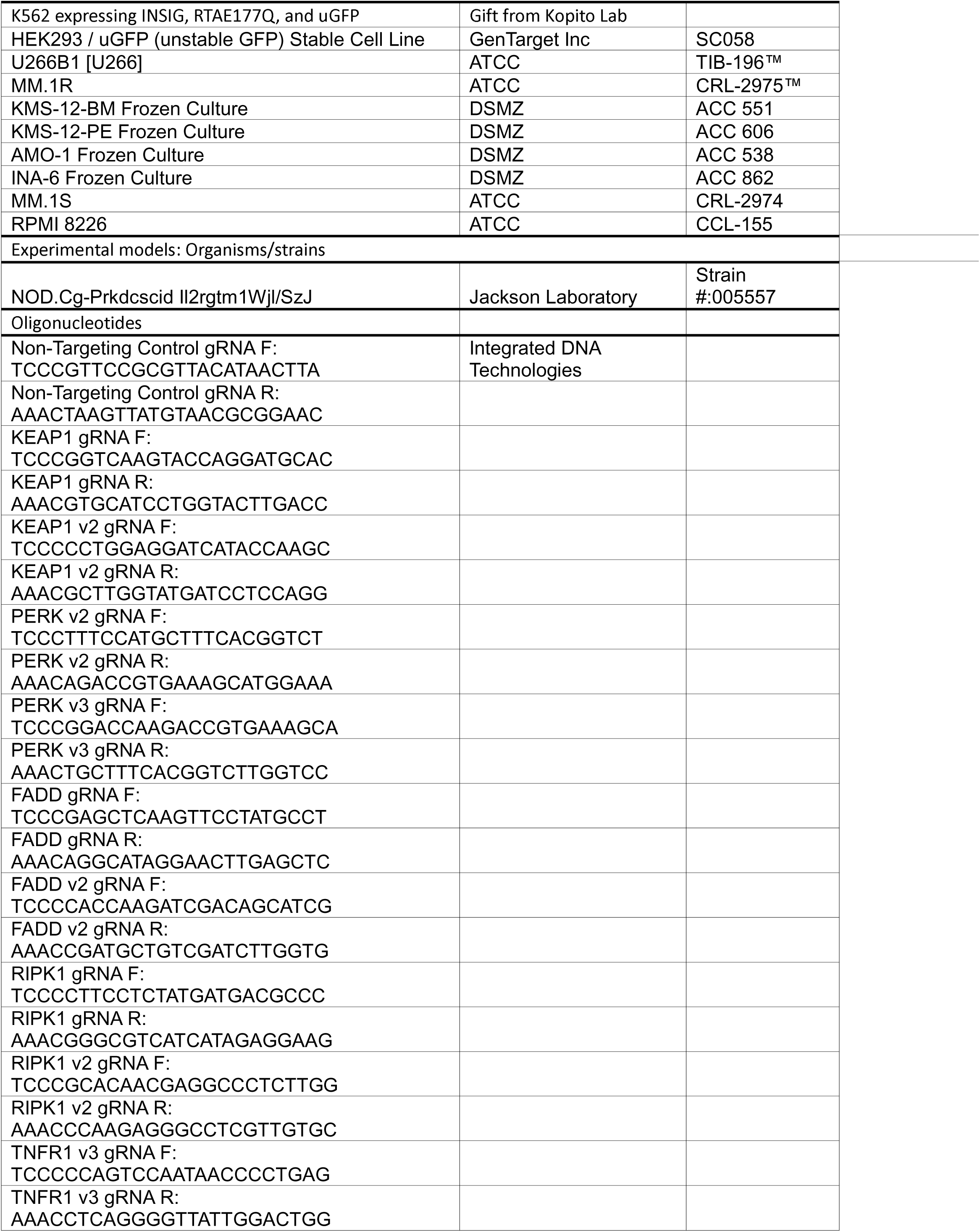
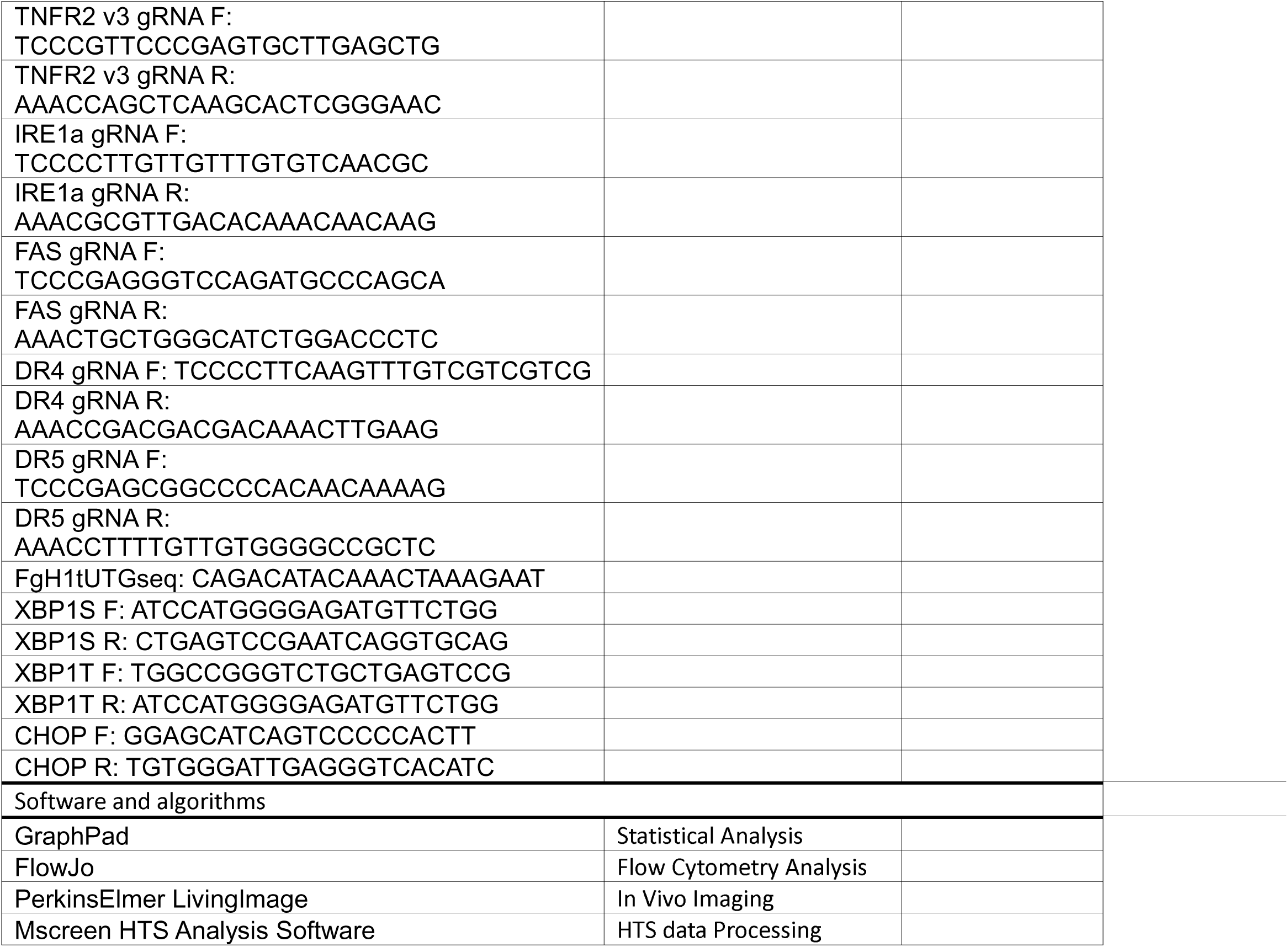

### Contact for Reagent and Resource Sharing

All inquiries for reagent or resource sharing should be directed to Qing Li at

### Experimental Models and Study Participants

#### Cell lines

MM.1S, RPMI8226, MM.1R, U266B1 were obtain from ATCC. KMS12-BM, AMO1, and INA-6 cells were obtained from DSMZ. NCIH929, OPM2, and KMS11 cells were a gift from the Talpaz lab. Cultures were routinely tested for mycoplasma contamination (Invivogen). MM.1S, RPMI8226, U226B1 were cultured in Roswell Park memorial Institute (RPMI) 1640 growth media with 10% Fetal Bovine Serum (FBS; Fisher) and 1x Penicillin-Streptomycin-Glutamine (Gibco); KMS12-BM, AMO1 and INA-6 were cultured as above but with 20% FBS. INA-6 was maintained in 10 ng/mL recombinant human IL-6 (Peprotech). K562 cells expressing INSIG-GFP, RTA^E177Q^-GFP, and uGFP were a gift from the Baldridge lab. Proteasome resistant AMO1 cell lines were developed by culturing with a gradual increase in BOR concentration as previously described^69,70^. RPMI8226 cells were transduced with Luciferase-2A-GFP Lentivirus (GenTarget Inc) according to manufacturer guidelines. RPMI8226-Luc cells were purified by fluorescence-activated cell sorting (SonyMA900) based on GFP expression. All cultured cells were maintained at 37°C with 5% CO_2_; cells were plated at 5e5 cells/mL unless specified.

#### Human Primary MM Samples

De-identified fresh bone marrow or peripheral blood samples were utilized from the University of Michigan hematologic malignancy repository bank at the University of Michigan, which are obtained and processed under the IRB approved studies HUM00002815 and HUM00066564. Samples were cultured with RPMI 1640 growth media with 20% Fetal Bovine Serum (FBS; Fisher), 10 ng/mL recombinant human IL-6 and 1x Penicillin-Streptomycin-Glutamine (Gibco) Xenograft Experiments: Animal studies were approved by the Institutional Animal Care and Use Committees at the University of Michigan. NOD.CG-Prkdc^scid^Il2rg^tm1Wjl^/SzJ (NSG) mice were obtained from The Jackson Laboratory and maintained in house. All animal studies were performed in accordance with IACUC guidelines. Male mice between 6-10 weeks of age were utilized for RPMI8226 xenograft transplant studies as previously described^49,50^. Mice received 175 cGy whole body irradiation (Kimtron Medical IC-320) followed by transplantation of 5e6 RPMI8226 cells with serial imaging and monitoring. All mice were maintained within the recommended tumor burden and survival endpoints per institutional regulations.

#### Plasmids

The GFP tagged null hong kong variant of alpha-1 antitrypsin (NHK) and transactivator plasmids were a gift from the Kopito lab. pLenti puro HA-Ubiquitin was a gift from Melina Fan (Addgene plasmid # 74218; http://n2t.net/addgene:74218; RRID:Addgene_74218). FgH1tUTG and FUCas9Cherry were a gift from Marco Herold (Addgene plasmid # 70183 and #70182; http://n2t.net/addgene:70183 and http://n2t.net/addgene:70182; RRIDs: Addgene70183 and 70182).

#### Lentivirus Generation & Transduction

Lentivirus was generated in 293T cells co-transfected with packaging plasmids pMD2.G and psPAX2 using FuGENE HD Transfection Reagent. CRISPR-CAS9 MM.1S cells were transduced by spinfection with lentivirus at 800 x g for 1.5h at 30 deg C followed by 1h incubation at 37C. K562 ells were transduced by infection with lentivirus at 800xg at RT for 1.75h.

### Method Details

#### Small Molecule Screening

NHK-GFP K562 cells were induced with doxycycline (0.75 µg/mL) for 16 hours. For screening, doxycycline was removed, and cells were plated on 384 well plates (Corning 3701) containing small molecules with 20 µM emetine in phenol red free RPMI 10% FBS. Compounds were pre-dispensed into 384 well plates using a Echo650 acoustic dispenser from Beckman Coulter. DMSO vehicle control was utilized with less than 0.5% DMSO. After four-hour incubation, cells were analyzed with automated flow cytometry (Biorad Ze5) with DAPI (1 µg/mL) dead cell exclusion. Primary hits were defined as >3 STDEV above DMSO control and were further prioritized as having >20% increase in MFI over DMSO control. Primary hits were validated in triplicate and were selected based on having at least three out of four instances with >3 STDEV above DMSO. Remaining compounds were then testes in 8-point concentration response curves in duplicate. Lead compounds with an IC_50_ of ≤20 µM were then tested in orthogonal assay to ensure target/mechanism specificity.

#### Small Molecule Treatment

*Selected lead* small molecule inhibitors were reordered/obtained commercially as described in the key resource table. For commercially acquired compounds, small molecules were dissolved in DMSO with a final concentration ≤0.1% DMSO. Emetine was dissolved in tissue culture grade H_2_O with a final concentration of ≤0.1%.

#### GFP Tagged ERAD Substrate Steady State Degradation

Expression of NHK-GFP, INSIG-GFP, RTA^E177Q^-GFP, and uGFP in K562 cells were induced with 0.75 µg/mL doxycycline for 16h. Doxycycline was removed, and K562 cells were plated with 20 µM emetine in phenol red free RPMI 10% FBS and small molecule inhibitors at specified concentrations for select timepoints between 0-21 hours. DAPI (1 µg/mL) was added for dead cell exclusion and cells were immediately analyzed by flow cytometry (BD LSRFortessa).

#### Ubiquitin Immunoprecipitation

K562 NHK-GFP cells expressing HA-Ubiquitin were induced with doxycycline (0.75 µg/mL) for 16 hours. Doxycycline was removed and cells were plated with 20 µM emetine with respective treatments. Following 4h treatment, cells were harvested and washed with PBS. Cells were lysed with IP Lysis buffer (Pierce) with cOmplete Mini Protease (Roche) and phosphatase inhibitor (Roche). Protein was quantified with the micro-BCA protein quantitation assay (Thermo Scientific). 500 µg total protein was loaded with HA-beads and immunoprecipation was performed with the HA-Tag IP/Co-IP kit per manufacturer’s instructions (Pierce) with 25 µL non-reducing sample buffer elution followed by the addition of β-mercaptoethanol. For immunoblot analysis 12.5 µL I.P and 10% of whole cell lysate input were loaded as a control.

#### Immunoblotting

Protein quantitation was performed with Qubit® Protein Assay (Thermo Fisher); 5-12 µg of protein was loaded per condition. Proteins were separated 4-20% Mini-Protean TGX gels (Biorad) with Tris/Glycine/SDS buffer (Biorad) and transferred to PVDF membranes. Total protein quantitation was performed with TotalStain Q (PVDF; Azure Biosystems) per manufacturer’s recommendations and membranes were blocked with 2% bovine serum albumin in Tris-buffered saline with Tween 20 (0.1%). Blots were assessed primary antibodies and secondary antibodies under conditions described in the key resources table.

#### Proteasome Activity

*MM.1S* cells were plated at 1.25e5 cells/mL and allowed to recover overnight. Cells were treated for 2h and immediately analyzed with the chymotrypsin-like proteasome Glo™ Cell Based Reagent per manufacturer’s instructions. Media only control was used as a blank and activity was normalized to DMSO control.

#### Cell Viability Assays

For MM cell line viability assays cells were plated at 1.25e5 cells/mL and allowed to recover for at least 2h prior to treatment. Following 12-72h treatment, ATP-dependent cell viability was measured with CellTiter Glo® (Promega) per standard manufacturer’s protocol and normalized to DMSO control. For viability measurements by Calcein AM (Biotium), cells were stained with 0.1 µM of Calcein AM for 30 minutes at room temperature (RT) after 24h, followed by the addition of DAPI (1 µg/mL) and flow cytometric analysis (Bio-Rad ZE5). Live cells were defined by Calcein AM^+^ and DAPI^-^ and normalized to the DMSO control. At 48 h treatment, Readyprobes ™ Cell Viability Imaging Kit (Invitrogen) was utilized per manufacturer’s instructions with flow cytometric analysis (Bio-Rad ZE5).

#### Annexin V Analysis of Early Apoptosis

At respective timepoints, 1.5e5 cells were collected and washed with Hanks’ Balanced Salt Solution (GIBCO) with 3% bovine calf serum (Cytiva). Cells were resuspended in Annexin Binding Buffer (Invitrogen) with Annexin V (5 µL) and DAPI (5 µg/mL) and incubated for 15 min at RT followed by immediate analysis by flow cytometry (BD LSRFortessa). Uniform gating based on forward and side scatter for single cell events was performed.

#### Protein Digestion and TMT labeling

MM.1S cells were treated with ERAD inhibitors for 4h in the presence of 50 uM emetine. Cells were harvested, washed with PBS at 4C, lysed with RIPA lysis buffer and protein was quantified with BCA assay. 100 µg total protein per condition were submitted to Proteomics Resource Facility at the University of Michigan for processing and mass spectrometry data acquisition. Briefly, upon reduction (5 mM DTT, for 30 min at 45 C) and alkylation (15 mM 2-chloroacetamide, for 30 min at room temperature) of cysteines in samples, the proteins were precipitated by adding 6 volumes of ice-cold acetone followed by overnight incubation at −20° C. The precipitate was spun down, and the pellet was allowed to air dry. The pellet was resuspended in 0.1M TEAB and overnight (∼16 h) digestion with trypsin/Lys-C mix (1:50 protease:protein (for solution digestion) at 37° C was performed with constant mixing using a thermomixer.

The TMT 16-plex reagents (ThermoFisher Scientific; A44521) were dissolved in 20 µl of anhydrous acetonitrile and labeling was performed by transferring the entire digest to TMT reagent vial and incubating at room temperature for 1 h. Reaction was quenched by adding 8 µl of 5% hydroxyl amine and further 15 min incubation. Labeled samples were mixed together, and dried using a vacufuge. An offline fractionation of the combined sample (∼300 µg) into 12 fractions was performed using high pH reversed-phase chromatography (Zorbax 300Extend-C18, 2.1mm x 150 mm column on an Agilent 1260 Infinity II HPLC system). Fractions were dried and reconstituted in 9 µl of 0.1% formic acid/2% acetonitrile in preparation for LC-MS/MS analysis. Samples were labeled with TMT mass tag channels as described in TMT Infor (Supplemental Table 1).

#### Liquid chromatography-mass spectrometry analysis

To obtain superior quantitation accuracy, we employed multinotch-MS3, which minimizes the reporter ion ratio distortion resulting from fragmentation of co-isolated peptides during MS analysis ^71^. Orbitrap Ascend Tribrid equipped with FAIMS source (Thermo Fisher Scientific) and Vanquish Neo UHPLC was used to acquire the data. Two µl of the sample was resolved on an Easy-Spray PepMap Neo column (75 µm i.d. x 50 cm; Thermo Scientific) at the flow-rate of 300 nl/min using 0.1% formic acid/acetonitrile gradient system (3-19% acetonitrile in 72 min;19--29% acetonitrile in 28 min; 29-41% in 20 min followed by 10 min column wash at 95% acetonitrile and re-equilibration) and directly spray onto the mass spectrometer using EasySpray source (Thermo Fisher Scientific). FAIMS source was operated in standard resolution mode, with a nitrogen gas flow of 4.2 L/min, and inner and outer electrode temperature of 100 °C and dispersion voltage or −5000 V. Two compensation voltages (CVs) of −45 and −65 V, 1.5 seconds per CV, were employed to select ions that enter the mass spectrometer for MS1 scan and MS/MS cycles.

Mass spectrometer was set to collect MS1 scan (Orbitrap; 400-1600 m/z; 120K resolution; AGC target of 100%; max IT in Auto) following which precursor ions with charge states of 2-6 were isolated by quadrupole mass filter at 0.7 m/z width and fragmented by collision induced dissociation in ion trap (NCE 30%; normalized AGC target of 100%; max IT 35 ms). For multinotch-MS3, top 10 precursors from each MS2 were fragmented by HCD followed by Orbitrap analysis (NCE 55; 45K resolution; normalized AGC target of 200%; max IT 200 ms, 100-500 m/z scan range).

#### qPCR

RNA was extracted using TRIzol Reagent (Invitrogen) following manufacturer guidelines. 6 uL of linear acrylamide was added after phase extraction to help precipitate RNA. RNA was reverse transcribed to cDNA using the High-Capacity cDNA Reverse Transcription Kit with Rnase Inhibitor (Applied Biosystems) following manufacturer guidelines. qPCR was performed (Applied Biosystems QuantStudio 3) using 10 ng of cDNA with SYBR Green PCR Master Mix (Applied Biosystems). qPCR primers are listed in Table 3.

#### Generation of gRNA Plasmids

gRNA sequences were designed using Benchling and cloned into the FgH1tUTG plasmid as previously described^72^. gRNA oligonucleotides are listed in Table 2. Successful cloning of gRNA sequence into FgH1tUTG was confirmed by Sanger sequencing (Azenta) using FgH1tUTGseq (5’-CAGACATACAAACTAAAGAAT-3’).

#### Generation of CRISPR-Cas9 KO Lines

MM.1S cells were transduced with FUCas9-Cherry. 72 hours after transduction, MM.1S cells were purified for Cas9 expression by fluorescence-activated cell sorting (SonyMA900) for mCherry expression to generate the MM.1S-Cas9 line. To generate MM.1S CRISPR KO lines, MM.1S-Cas9 cells were transduced with the generated gRNA plasmids. Cells were used for experiments at least 3-7 days after gRNA expression was induced by doxycycline (1 µg/mL). Confirmation of KO was completed by immunoblotting.

#### Caspase Activity Assay

For caspase 8 activity assay, cells were treated for specified timepoints, at which time CaspaTag Caspase 8 Fluorescein reagent was added. Cells were incubated for 15 min at 37C under 5% CO2, at which time cells were harvested, washed with 1 mL Annexin Binding buffer and resuspended with Annexin V APC (5uL) and DAPI (5 µg/mL). 15 min at RT followed by immediate analysis by flow cytometry (BD LSRFortessa). Caspase 3/7 activity was measured with Cell Event Caspase-3/7 (Invitrogen) per manufacturer’s instructions with DAPI (1 µg/mL) dead cell exclusion. Caspase 3/7 activity was quantified by flow cytometry analysis (BD LSRFortessa) at specified timepoints.

#### Immunofluorescence

5e5 MM.1S cells were plated in 1 mL media and treated with ERAD inhibitors for 2 hours. Cells were collected and stained in 1 mL recombinant cholera toxin Subunit B Alexa Fluor ™ 555 (1 µ/mL), Annexin V FITC, and NucRed™ Live 647 ReadyProbes™ (2 drops) Reagent in complete media for 15 min at 4C. Cells were washed, fixed with 4% formaldehyde for 10 min at RT, and mounted with Prolong™ Gold Antifade Reagent. Fluorescence was imaged on a THUNDER Imaging System (Leica) using a 100X objective.

#### ROS Measurement

MM.1S cells were loaded with 10 µM CM-H2DCFDA (Diluted 1:500 from 5 mM DMSO stock) in prewarmed HBSS for 30 min at 37C. Cells were washed with HBSS and resuspended in standard media under specified treatment conditions. After 2 hours, 200 uL of cells were collected, followed by the addition of DAPI (5 µg/mL) and flow cytometric analysis (Bio-Rad ZE5). For positive control, 150 µM H2O2 was added for 1h.

#### Cell Surface Immunophenotyping

At specified timepoints, 1.5e5 cells were collected and incubated with Human TruStain FcX™ (Biolegend) for 5 min at 4C, followed by the addition of cell surface antibodies for 10 min at 4C. Cells were washed and resuspended in Annexin Binding Buffer (Invitrogen) with Annexin V (5 µL) and DAPI (5 µg/mL). Samples were incubated for 15 min at RT followed by immediate analysis by flow cytometry (BD LSRFortessa).

#### Xenograft Transplant Imaging

Starting from week 3 post-transplant, mice were administered RTA408 (5 mg/kg) or vehicle (10% DMSO in Corn Oil) intraperitoneally every other day until takedown. Bioluminescence imaging (Perkin Elmer IVIS Spectrum) was conducted every week starting 3 weeks post-transplant until takedown with intraperitoneal luciferin.

### Quantitation and Statistical Analysis

#### Proteomic Analysis

Proteome Discoverer (v3.0; Thermo Fisher) was used for data analysis. MS2 spectra were searched against SwissProt human protein database (v2023-09-13) using the following search parameters: MS1 and MS2 tolerance were set to 10 ppm and 0.6 Da, respectively; carbamidomethylation of cysteines (57.02146 Da) and TMT labeling of lysine and N-termini of peptides (304.2071 Da) were considered static modifications; oxidation of methionine (15.9949 Da) and deamidation of asparagine and glutamine (0.98401 Da) were considered variable. Identified proteins and peptides were filtered to retain only those that passed ≤1% FDR threshold. Quantitation was performed using high-quality MS3 spectra.

Z’ calculated as previously described^73^. IC_50_ was determined by variable slope-four parameter dose response curve fits. Statistical analyses used in these studies include t tests (2 samples), one-way ANOVA (>2 samples) or two-way anova (>2 samples with two parameters) in GraphPad Prism and are specified in the corresponding figure legends. *p≤0.05, ** p≤0.01, ***p≤0.001, ****p≤0.0001.

## Data Availability

This study did not generate unique reagents. The mass spectrometry proteomics data have been deposited to the ProteomeXchange Consortium via the PRIDE^74^ partner repository with the dataset identifier PXD061058“

## Author Contributions

E.M.K and Q.L conceptualized, designed the studies, and wrote the manuscript. E.M.K, S.M., O.W., A.M.R, M.K., J.G. performed experiments and data analysis. A.A., M.T., M.P, and Q.L. supervised and aided in experimental design. All authors performed manuscript review and editing. Funding acquisition was obtained by Q.L.

## Acknowledgements

We would like to thank patients and their families for their biospecimen contributions. We thank the Kopito and Baldridge labs for their gift of K562 cell lines and plasmids. We thank Ursula Jakob for her assistance with lipid raft microscopy. We thank the protein resource facility for their assistance with quantitative proteomics. E.M.K was supported by the Research Training Award for Fellows from the American Society of Hematology and the pre-Career Development Award from the National Oncology Program. These studies were funded by Michigan Drug Discovery, Protein Folding Disease Initiative, Rogel Cancer Center Discovery Award, R01HL150707, and R01HL174566. The graphical abstract and schematic of the extrinsic apoptotic pathway were generated with biorender.com.

## Conflicts of Interests

E.M.K, S.M., O.Y.W., A.M.R., M.K, J.L.G., B.J.O, A.A., M.T. Q.L. have no relevant competing interests to declare. M.P. receives honoraria/consulting from Janssen/Johnson & Jonson, Bristol Myers Squibb, Sanofi, GlaxoSmithKline, Pfizer and Research Funding (To Institution): AbbVie, AstraZeneca, Bristol Myers Squibb, Janssen, Pfizer, Sanofi, Regeneron.

During the preparation of this work the author(s) used ChatGTP in order to proofread the discussion. After using this tool/service, the author(s) reviewed and edited the content as needed and take(s) full responsibility for the content of the publication.

---

**Supplemental Figure 1:**
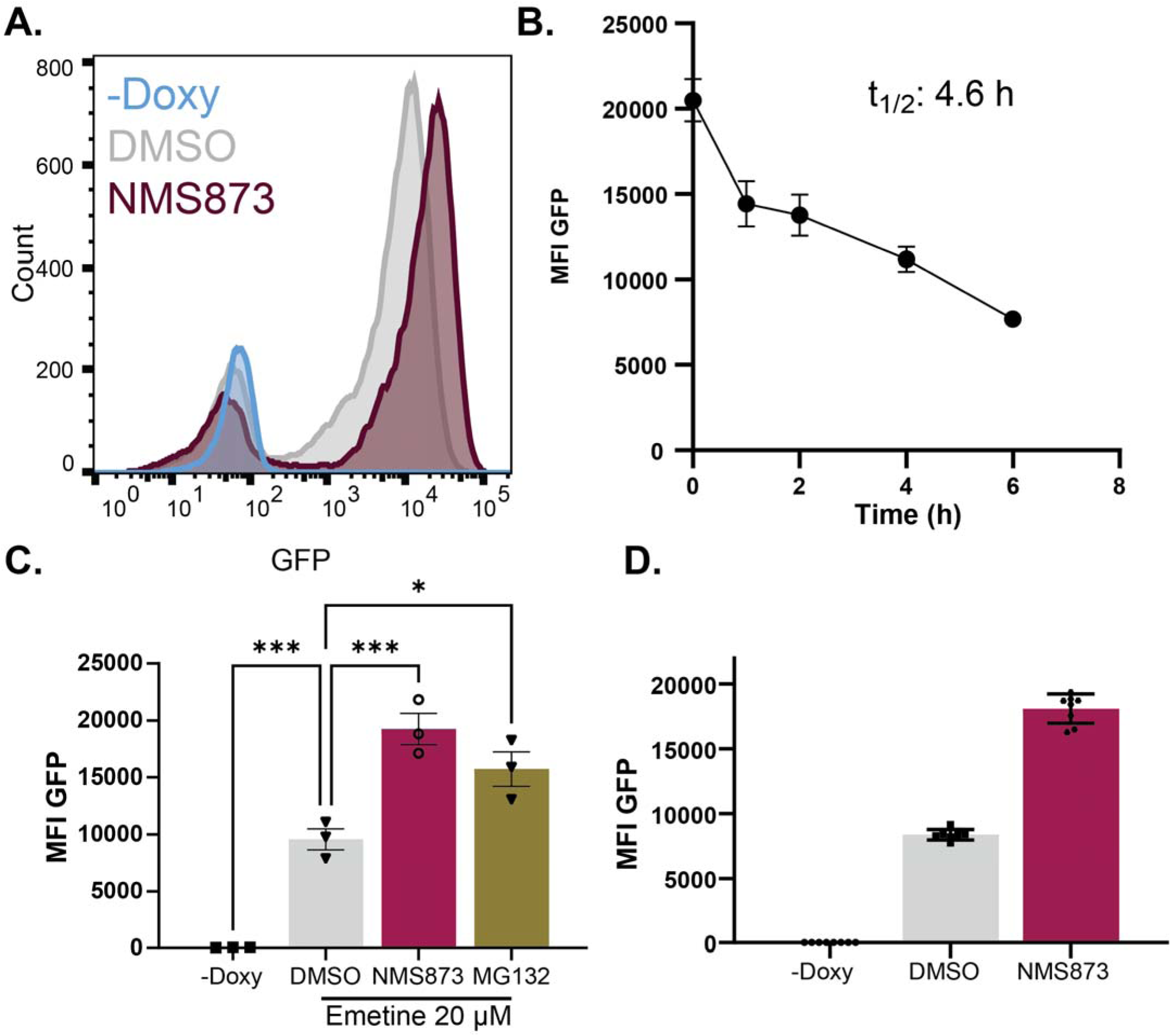
Validation of Screening Approach for ERAD Substrate Degradation. A. Representative flow cytometry plot for mean fluorescence of NHK-GFP in K562 cells treated with 20 µM emetine and DMSO or 10 µM NMS873 for 4h. -doxycyline is a negative control. B. Flow cytometry analysis of NHK-GFP degradation in live K562 cells in the presence of 20 µM emetine between 0-6h. N=3 C. Quantitation of mean fluorescence intensity of steady state degradation for NHK-GFP in K562 with DMSO or 10 µM NMS873 or MG132 at 4h. N=3 D. C. Quantitation of mean fluorescence intensity of steady state degradation for NHK-GFP in K562 with DMSO or 10 µM NMS873 at 4h by automated flow cytometry in a 384 well plate used to calculate Z’. N=16-32 technical replicates. Mean±STDEV. Statistical analysis performed with a one-way ANOVA with Dunnett’s multiple comparison test. *p≤0.05, ***p≤0.001

**Supplemental Figure 2:**
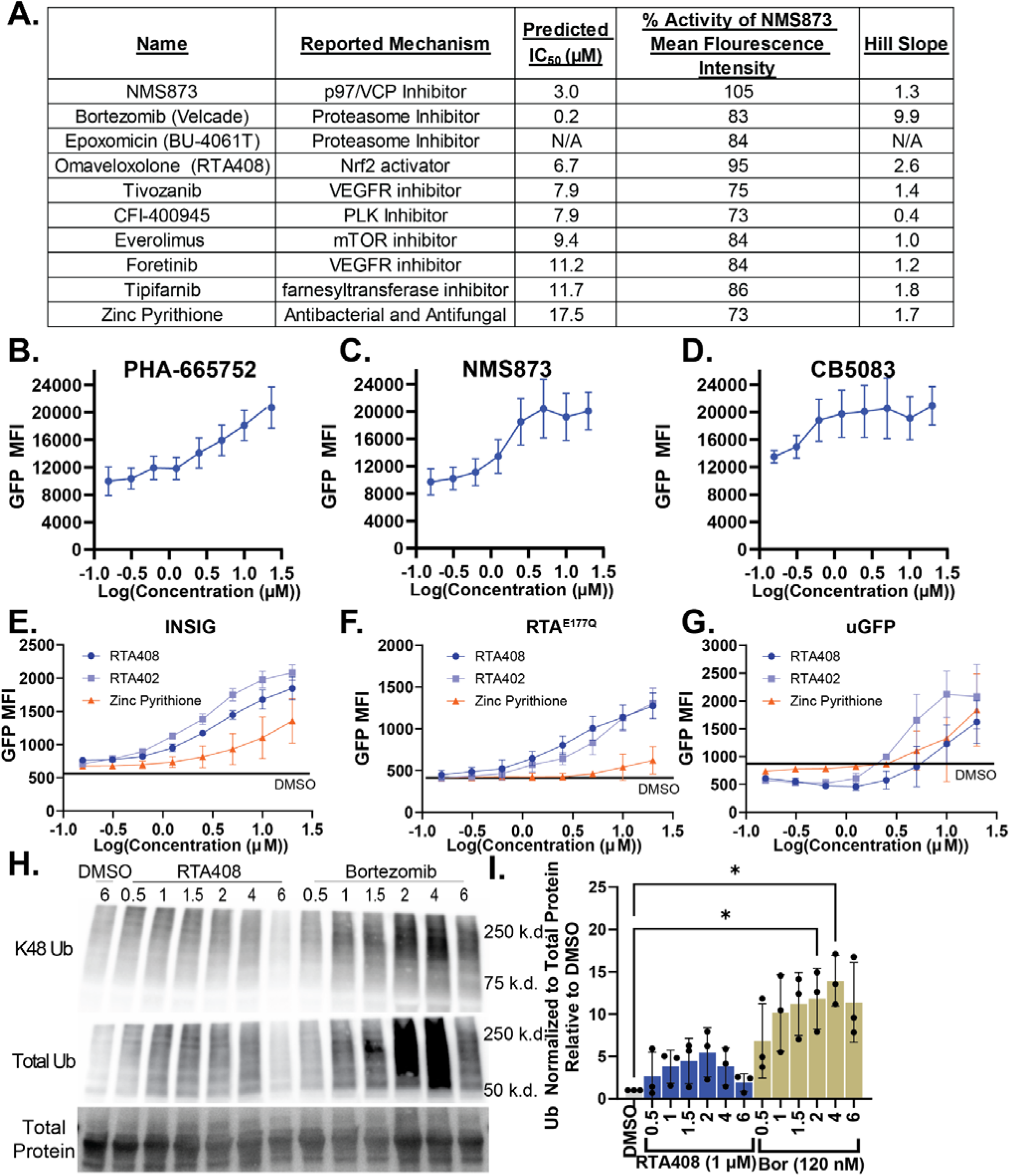
Validation of FDA Repurposing Library Hits. A. Table summarizing the top 10 hits from the FDA repurposing library with IC_50_, relative activity of NMS873 control, and hillslope estimated by variable slope-four parameter dose response curve based on curve represented in Figure 1A. B-D. Dose response curve for inhibition of NHK-GFP steady state degradation by PHA-665752(B), NMS873 (C), or CB5083 (E) at 156 nM-20 µM at 4h in K562 cells. E-G. Steady state degradation INSIG-GFP (E), RTA^E177Q^-GFP (F), and uGFP (G) with DMSO or 156 nM-20 µM RTA408, RTA402, or Zinc Pyrithione at 1h in K562 cells. H. Representative immunoblot of K48, total ubiquitin and total protein quantitation in MM.1S cells following treatment with DMSO, 1 µM RTA408, 10 µM NMS873, or 120 nM BOR for 0.5-6 h. I. Relative quantitation of total ubiquitin normalized for total protein quantitation from immunoblots in H. N=3. Mean±STDEV. Statistical analysis performed with a Kruskal-Wallis test with Dunn’s multiple comparisons. *p≤0.05.

**Supplemental Figure 3.**
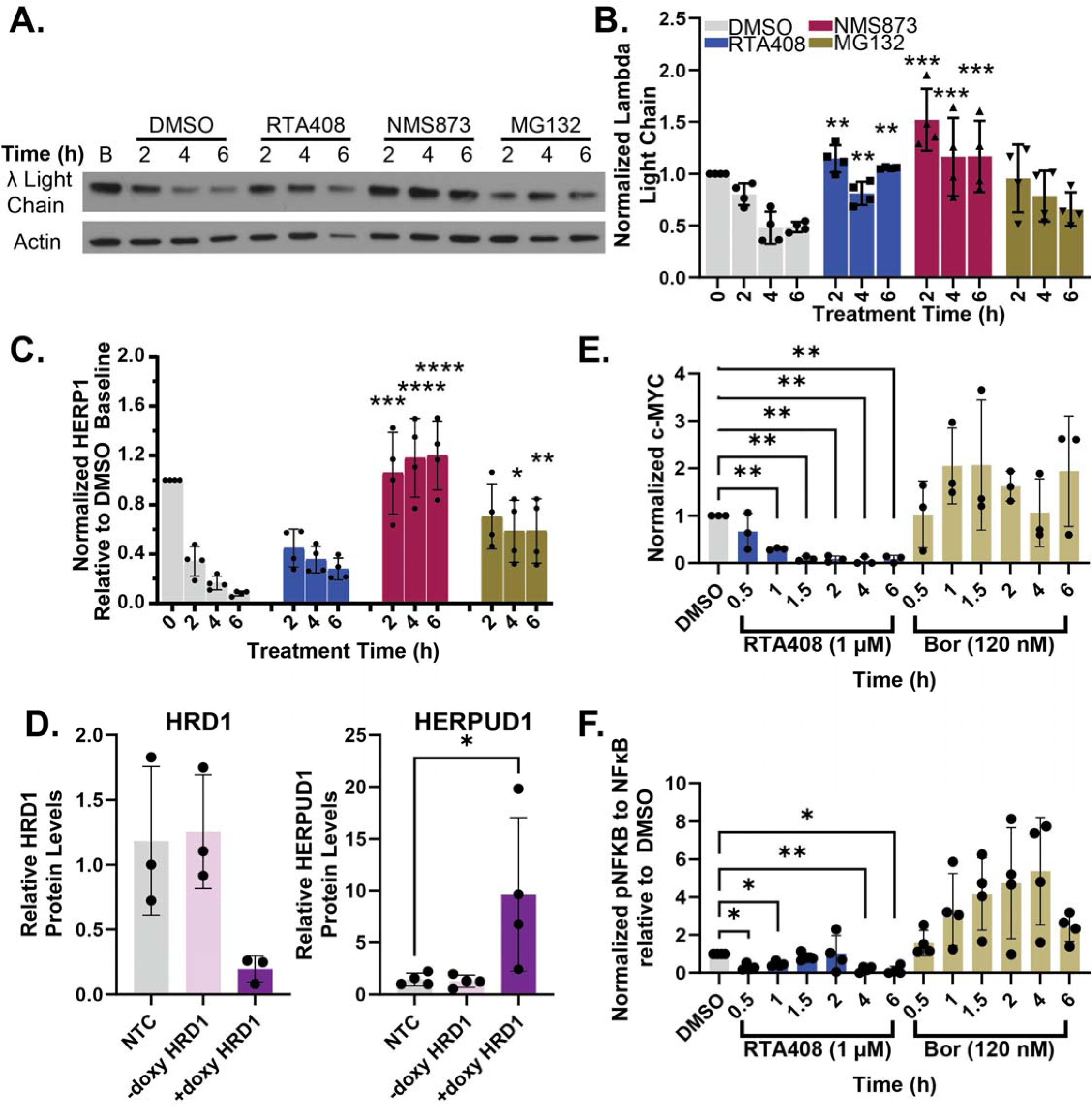
Validation of Proteomic Analysis. A-B. Immunoblot (A) and relative quantitation (B) for steady state degradation (50 µM emetine) with DMSO, 1 µM RTA408, 10 µM NMS873, or 10 µM MG132 in MM.1S. B-E. Relative quantitation for immunoblot analysis of steady state degradation of HERPUD1 (C), HERPUD1 in HRD1 KO MM.1s (D), c-MYC(E), or pNFKB(F). Statistical analysis with a two-way ANOVA with Tukey’s multiple comparisons (B-C; DMSO 0h was excluded from statistical analysis) or one-way ANOVA with Dunnett’s multiple comparisons (D-F). * p≤0.05, **p≤0.01, ***p≤0.001, **** p≤0.0001

**Supplemental Figure 4.**
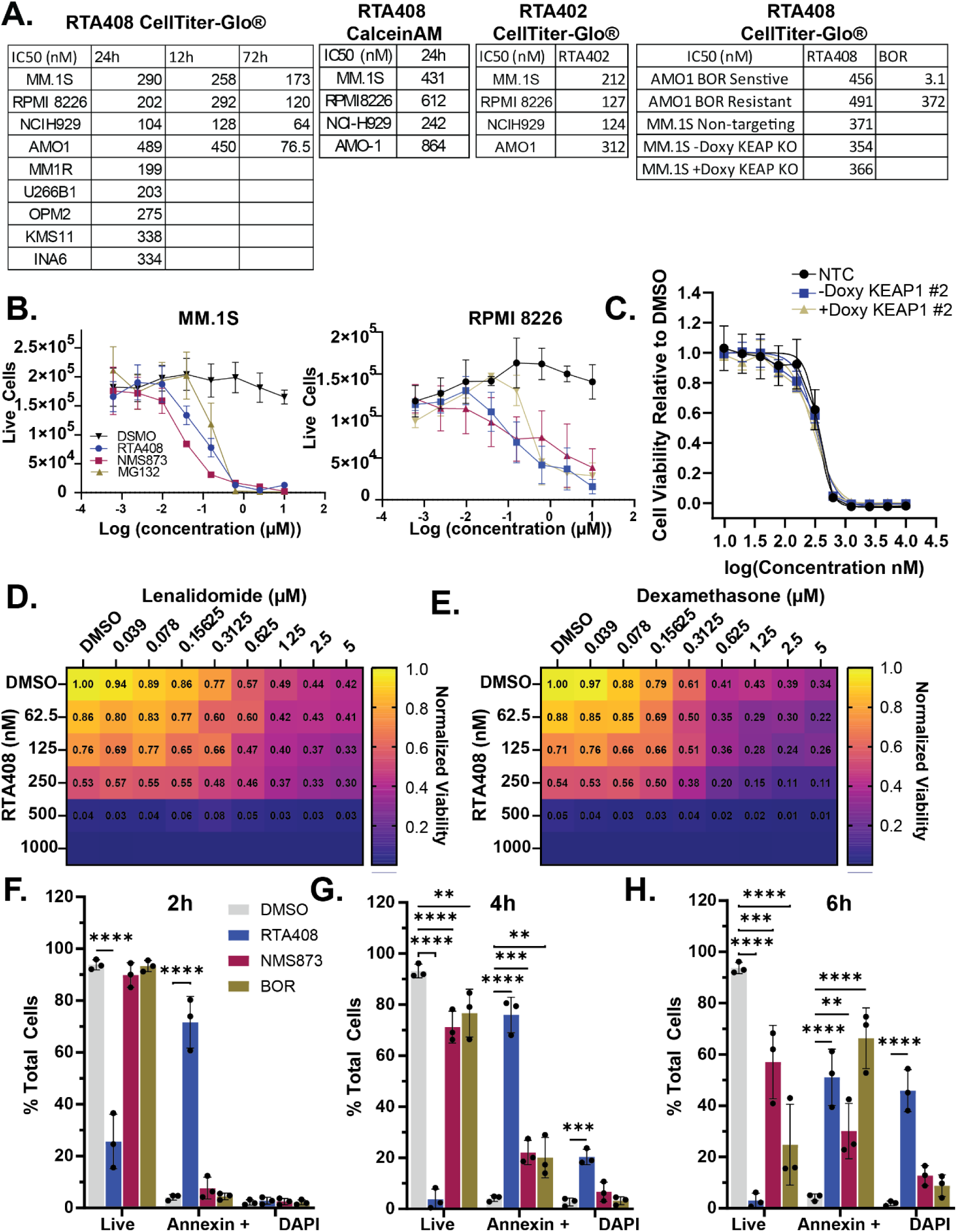
RTA408 cytotoxicity in MM cells. A. Summary of IC_50_ for RTA408 cytotoxicity between 12-72h determined by CellTiter-Glo or Calcein AM staining. B. Cell viability determined by live/dead staining (Invitrogen) following 48 h treatment with 10 nM-10 µM RTA408, NMS873, or MG132 in MM.1S and RPMI8226 cells. C-D. Heatmap with viability measured by CellTiter-Glo® in MM.1S cells 24h following 39nM-5 µM lenalidomide (C) or 39nM-5 µM dexamethasone (D) and RTA408 62.5-1000 nM cotreatment at 72h. Viability is normalized to DMSO control. F-H. Quantitation of live (AnnexinV-DAPI-), Annexin+ (AnnexinV+DAPI-) or dead (AnnexinV+DAPI+) population at 2(F), 4 (G), or 6h (H) in MM.1S cells. Statistical Analysis by 2-way ANOVA with Dunnett’s multiple comparison tests. **p≤0.01, ***p≤0.001, **** p≤0.0001.

**Supplemental Figure 5.**
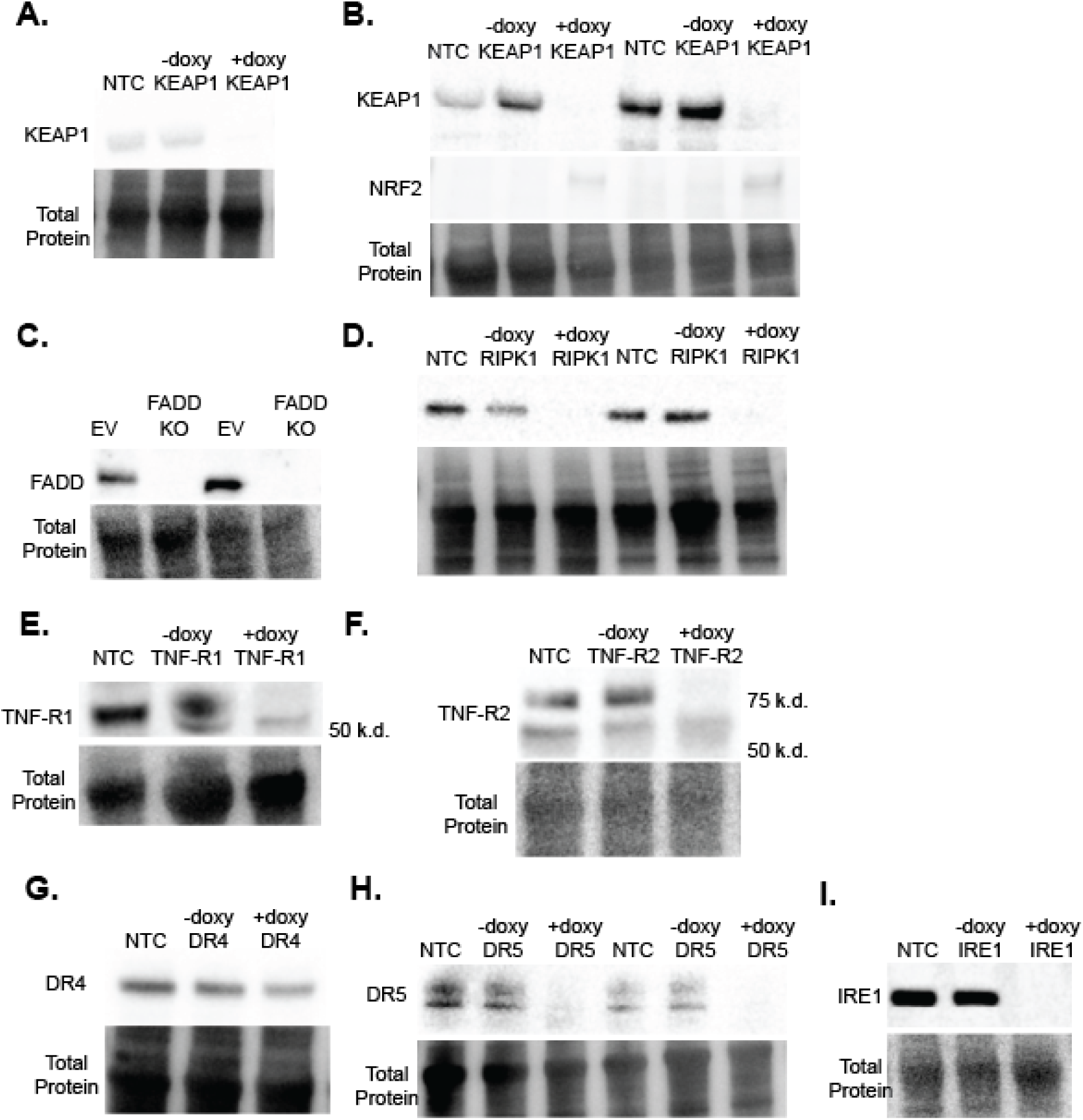
Validation of CRISPR-CAS9 Inducible KO. Representative immunoblot of NTC, and + or – doxycycline with sgKEAP1 #1 (A) and sgKEAP1 KO #2 (B), sgFADD (C), sgRIPK1(D), sgTNF-R1 (E), sgTNF-R2 (F), sgDR4 (G) or sgDR5 (H), or sgIRE1α (I) in MM.1S cells.

**Supplemental Figure 6.**
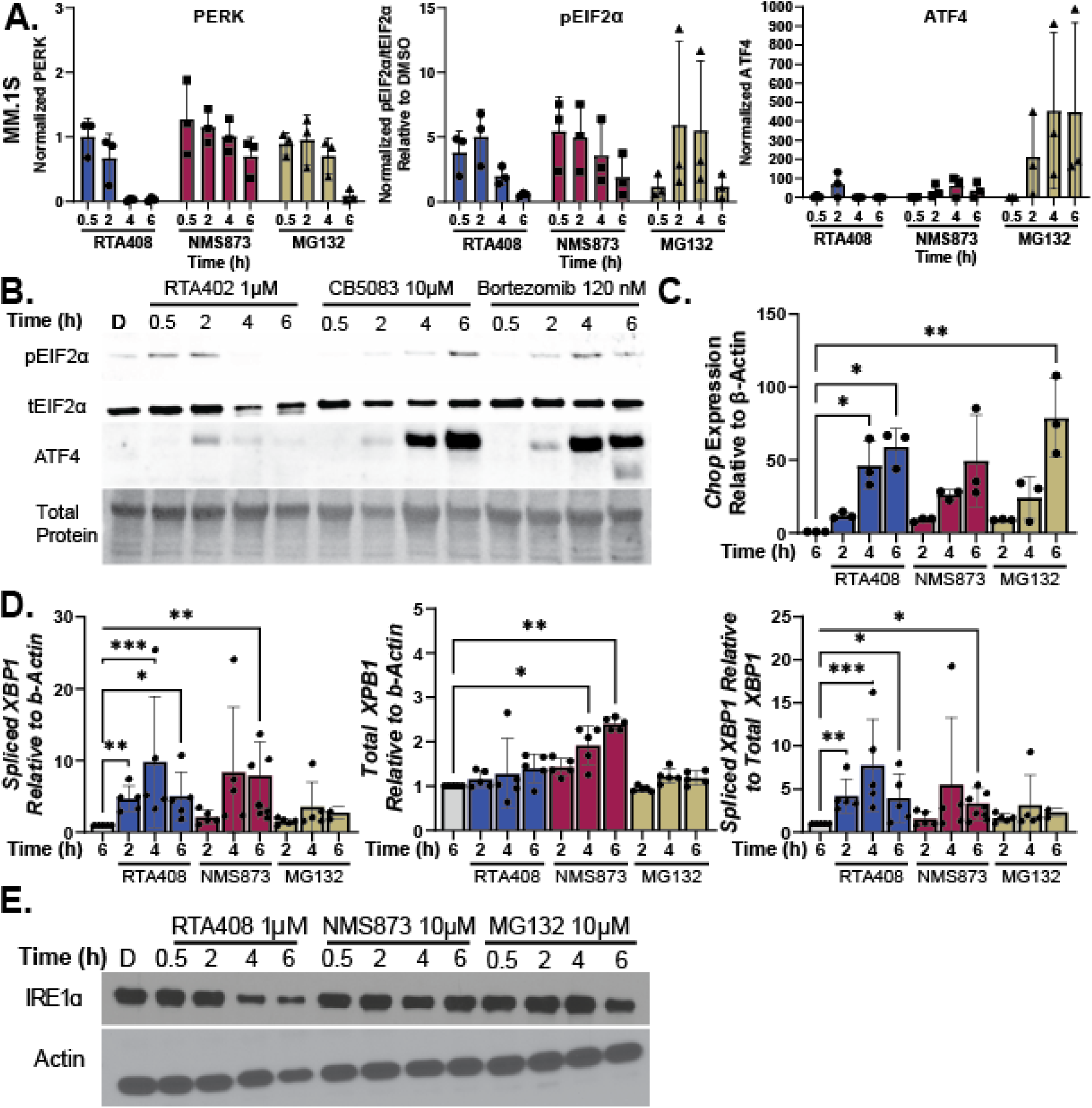
UPR Activation in MM.1S. A. Relative quantitation of PERK, pEIF2A, and ATF4 immunoblots for MM.1S treated with DMSO, 1 µM RTA408, 10 µM NMS873, or 10 µM MG132 in MM.1S for 0.5-6h. B. Immunoblot of pEIF2a, total EIF2a, PERK, ATF4, and β-Actin in MM.1s treated with 1 µM RTA402, 10 µM CB5083, or 120 nM BOR for 0.5-6h. C-D. Relative qPCR quantitation of CHOP (C), spliced or total XBP1 (D) relative to β-Actin in MM.1S treated with 1 µM RTA408, 10 µM NMS873, or 10 µM MG132 for 2-6h normalized to DMSO 6h control. E. Immunoblot of IRE1α, and β-Actin in MM.1s treated with 1 µM RTA408, 10 µM NMS873, or 10 µM MG132 for 0.5-6h. N=3-4. Mean±STDEV. No statistical analysis performed (A), statistical analysis with Kruskal-Wallis test with Dunn’s multiple comparisons test (C-D). *p≤0.05, **p≤0.01, ***p≤0.001

**Supplemental Figure 7.**
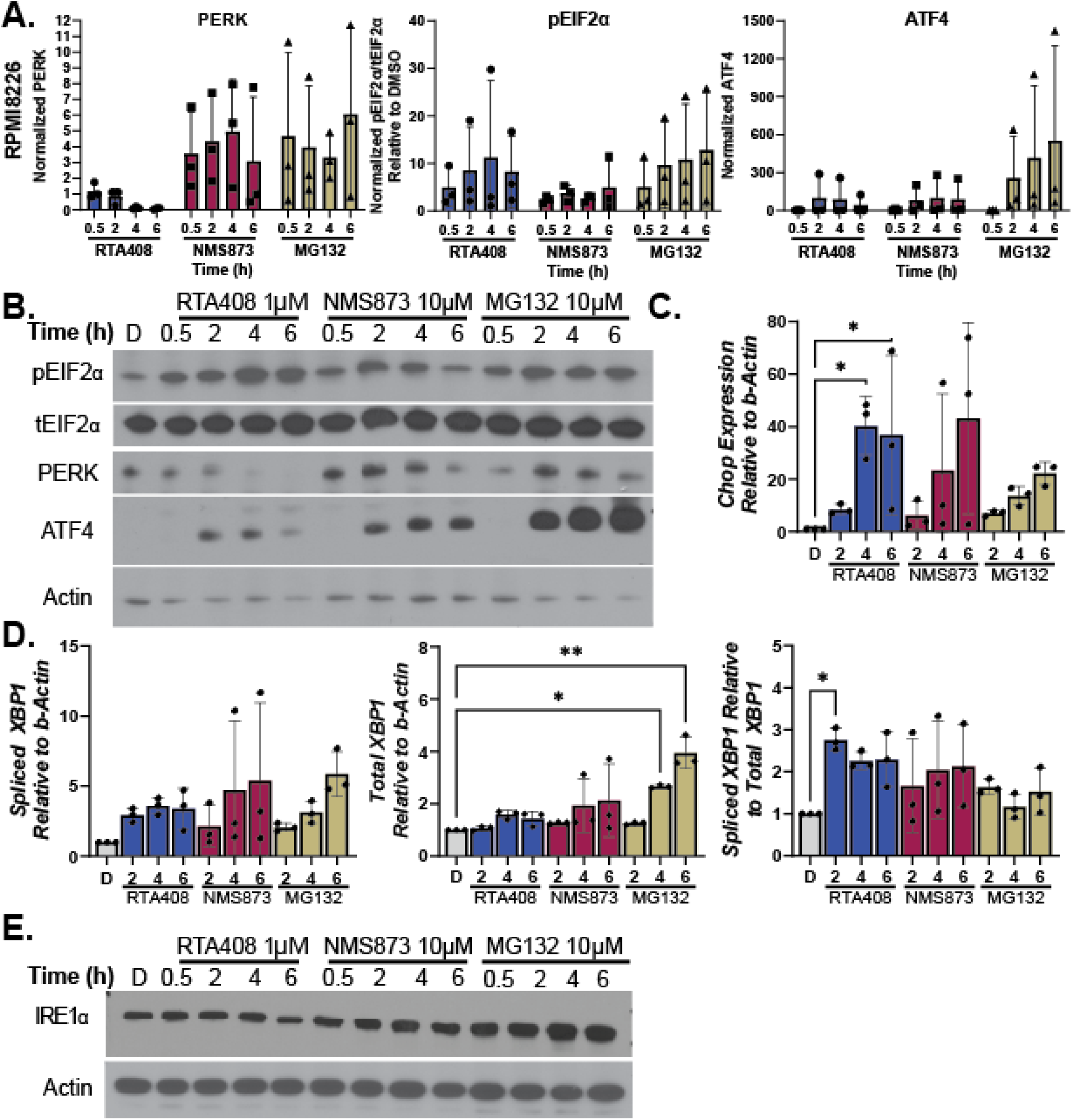
UPR Activation in RPMI8226. A-B. Relative quantitation (A) and Representative immunoblots (B) of PERK, pEIF2A, and ATF4 immunoblots for RPMI8226 treated with DMSO, 1 µM RTA408, 10 µM NMS873, or 10 µM MG132 for 0.5-6h. C-D. Relative qPCR quantitation of CHOP (C), spliced or total XBP1 (D) relative to β-Actin in RPMI8226 treated with 1 µM RTA408, 10 µM NMS873, or 10 µM MG132 for 2-6h normalized to DMSO 6h control. E. Immunoblot of IRE1α, and β-Actin in RPMI8226 treated with 1 µM RTA408, 10 µM NMS873, or 10 µM MG132 for 0.5-6h. N=3-4. Mean±STDEV. No statistical analysis performed (A), statistical analysis with Kruskal-Wallis test with Dunn’s multiple comparisons test (C-D)*p≤0.05, **p≤0.01

**Supplemental Figure 8.**
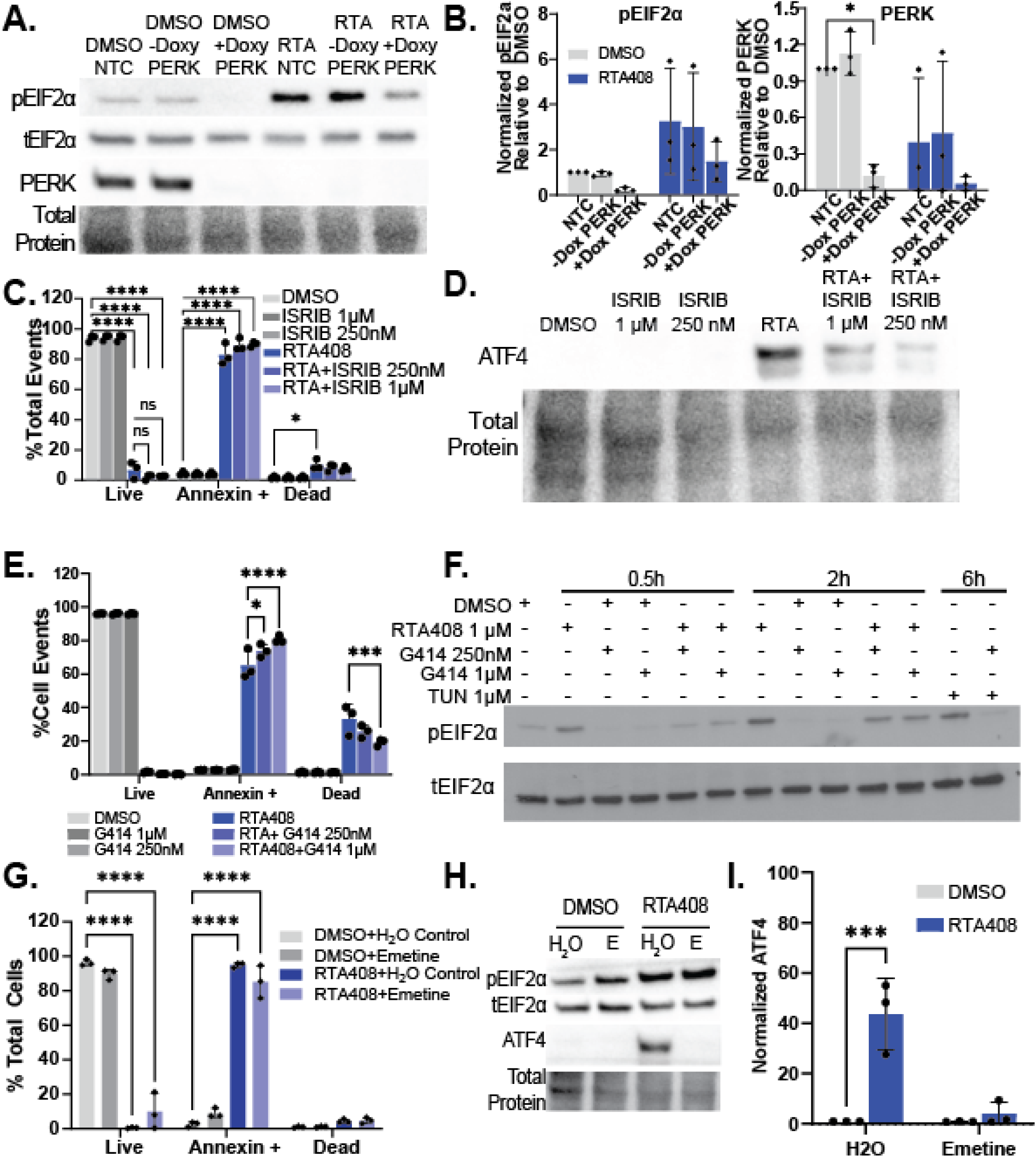
PERK Inhibition with ERAD Inhibition. A-B. Immunoblot (A) and relative quantitation (B) for pEIF2a, total EIF2a, PERK, and total protein in in MM.1S cells transduced with non-targeting control, or doxycycline inducible PERK KO (+or-doxycycline) treated with RTA408 1 µM for 2h. C and E. Quantitation of live (AnnexinV-DAPI-), Annexin+ (AnnexinV+DAPI-) or dead (AnnexinV+DAPI+) populations by flow cytometry in MM.1S cells treated with DMSO, RTA408 1 µM, 250 nM or 1 µM ISRIB (C) or GSK2606414 (G414) (E) for 4h. D. Representative immunoblot of ATF4 and total protein in MM.1S treated with with DMSO, RTA408 1 µM, 250 nM or 1 µM ISRIB for 2h. F. Representative immunoblot of pEIF2a and tEIF2a in MM.1S treated with with DMSO, RTA408 1 µM, 250 nM or 1 µM GSK2606414 (G414) for 0.5-2h. 1 µM Tunicamycin (TUN) at 6h was used as a positive control for G414 mediated inhibition of PERK signaling. G. Quantitation of Annexin V staining by flow cytometry in MM.1S cells treated with DMSO, RTA408 1 µM and H2O control or 50 µM emetine (labelled E) for 4h. H-I. Representative immunoblot (H) and relative quantitation (I) of pEIF2a, total EIF2a, ATF4, and total protein in in MM.1S at 2h with DMSO or RTA408 1 µM combined with H2O control or 50 µM emetine. N=3-4. Mean±STDEV.*p≤0.05, ***p≤0.001, ****p≤0.0001.

**Supplemental Figure 9:**
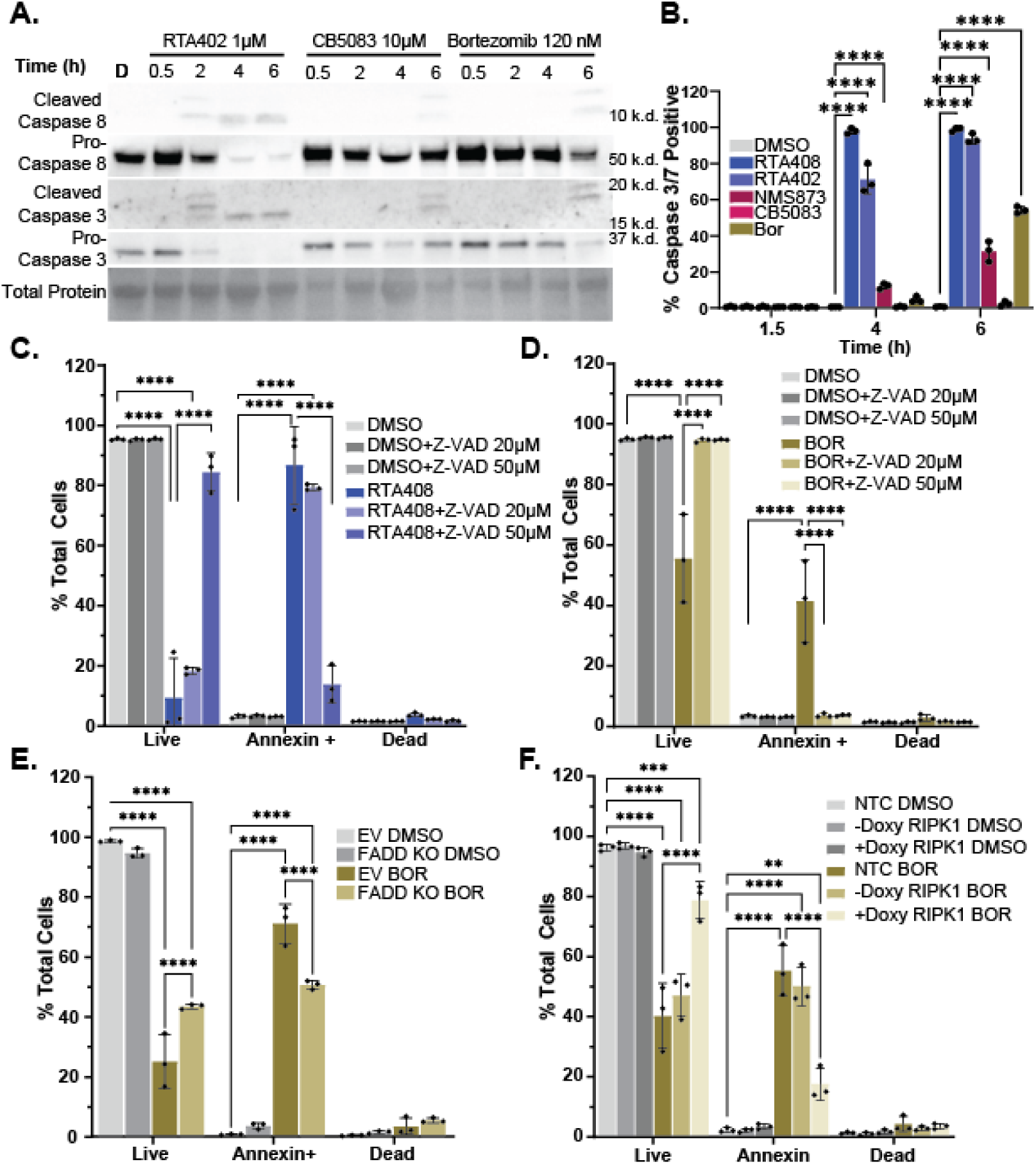
Pro-Apoptotic Signaling with ERAD Inhibition in MM.1S. A. Immunoblot of caspase 8 and 3 (cleaved and pro-forms) and total protein in MM.1s treated with DMSO, 1 µM RTA402, 10 µM CB5083 or 120 nM BOR for 0.5-6h.. B. Flow cytometry quantitation of caspase 3/7 activity in live MM.1S following DMSO, 1 µM RTA408 or RTA402, 10 µM CB5083 or NMS873, or 120 nM BOR for 1.5-6h. DMSO, RTA408, and BOR data are also represented in figure 5C. C-D. Flow cytometry quantitation of live (AnnexinV-DAPI-), Annexin+ (AnnexinV+DAPI-) or dead (AnnexinV+DAPI+) in MM.1S treated with 20-50 µM Z-VAD-FMK (C-D) and DMSO, 1 µM RTA408 for 4h (C), or 120 nM BOR for 6h (D). E-F. Flow cytometry annexin V staining in MM.1S cells transduced with empty vector (EV), non-targeting control (NTC), or doxycycline inducible FADD KO (E) or RIPK1 KO (F) treated with DMSO or 120 nM BOR for 6h. N=3. Mean±STDEV. Statistical analysis performed with a two-way ANOVA with Tukey’s multiple comparison test. ****p≤0.0001

**Supplemental Figure 10:**
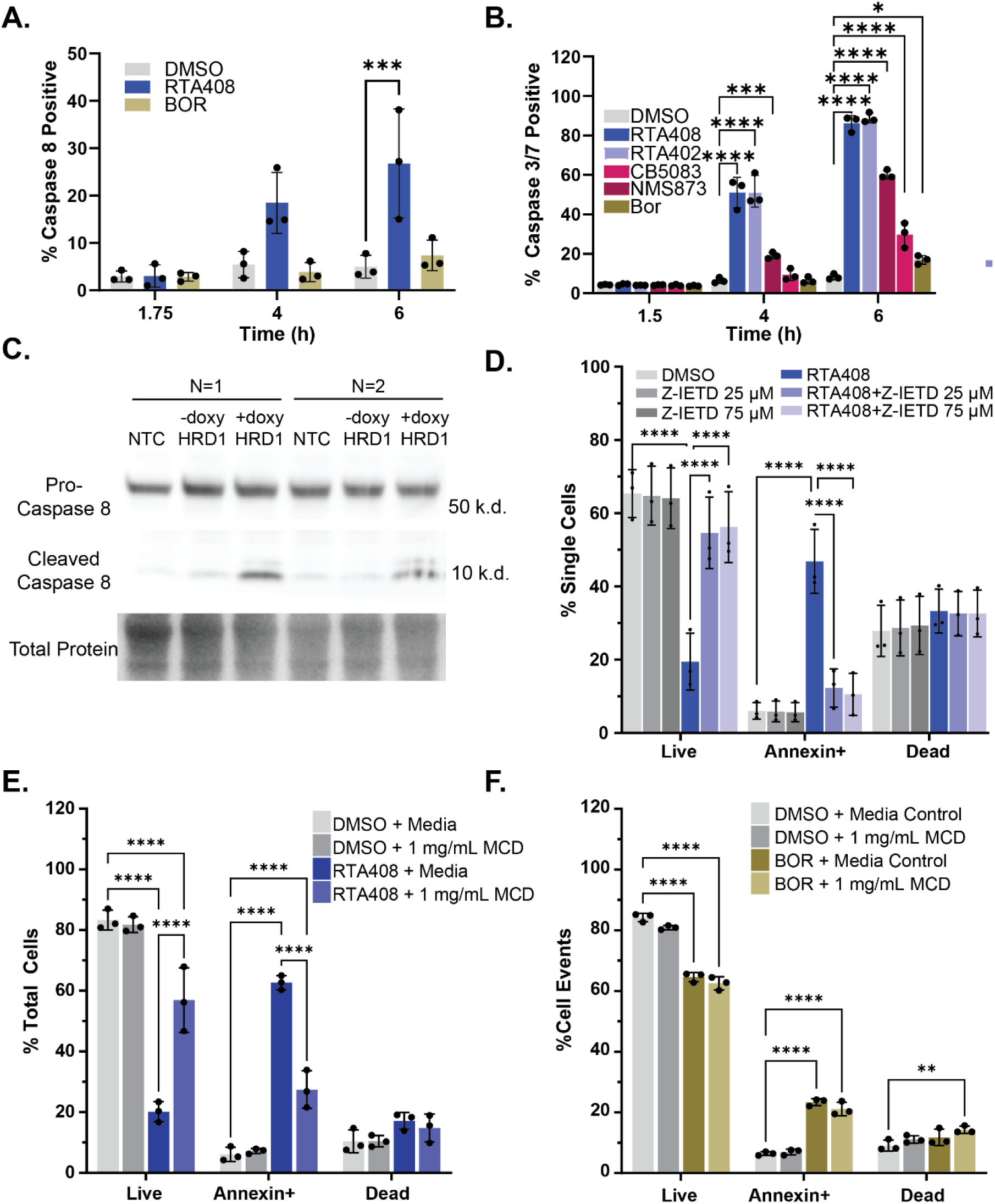
Pro-Apoptotic Signaling with ERAD Inhibition. A-B. Flow cytometry quantitation of caspase 8 (A) or caspase 3/7 (B) activity in live RPMI8226 following DMSO, 1 µM RTA408 or RTA402, 10 µM CB5083 or NMS873, or 120 nM BOR for 1.5-6h. C. Representative immunoblot of pro-caspase 8, cleaved caspase 8, and total protein quantitation in MM.1S transduced with non-targeting control (NTC) and doxycycline inducible HRD1 CRISPR-CAS9 KO (+ or – doxycycline). D. Flow cytometry quantitation of live (AnnexinV-DAPI-), Annexin+ (AnnexinV+DAPI-) or dead (AnnexinV+DAPI+) in RPMI8226 treated with 25-75 µM Z-IETD-FMK and DMSO or 1 µM RTA408 4h. E-F. Flow cytometry quantitation of Annexin V staining in RPMI8226 treated with 1 mg/mL MCD or media control and DMSO, 1 µM RTA408 for 4h (E) or 120 nM BOR for 6h (F). N=3. Mean±STDEV. Statistical analysis performed with a two-way ANOVA with Tukey’s multiple comparison test. *p p≤0.05, **p≤0.01, ****p≤0.0001

**Supplemental Figure 11:**
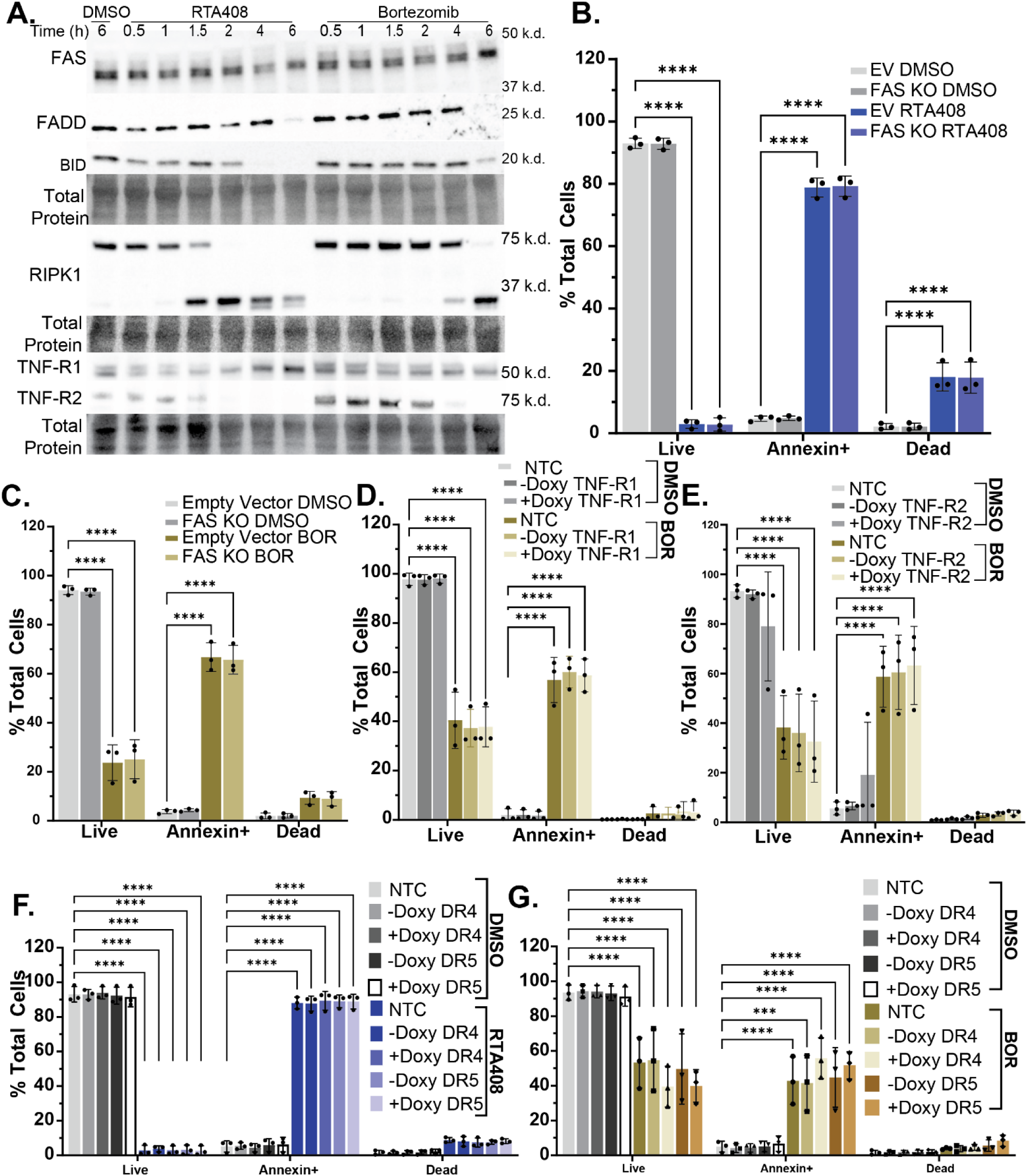
Investigation of Cell Death Receptor Mediated Activation of Caspase 8. A. Immunoblot of FAS, FADD, BID, RIPK1, TNF-R1, TNF-R2 and total protein levels in MM.1S treated with DMSO, 1 µM RTA408, or 120 nM BOR for 0.5-6h. B. Flow cytometry quantitation of live (AnnexinV-DAPI-), Annexin+ (AnnexinV+DAPI-) or dead (AnnexinV+DAPI+) MM.1S cells transduced with non-targeting control, or doxycycline inducible FAS treated with DMSO or 1 µM RTA408 for 4h. C-E. Flow cytometry quantitation of AnnexinV staining in MM.1S cells transduced with non-targeting control, or doxycycline inducible FAS (C), TNF-R1 (D), TNF-R2 (E), (+or-doxy) treated with DMSO or 120 nM BOR for 6h. F-G. Flow cytometry quantitation of Annexin V staining in MM.1S cells transduced with non-targeting control, or doxycycline inducible DR4 or DR5 KO (+or-doxy) treated with DMSO or 1 µM RTA408 for 4h (F) or DMSO or 120 nM BOR for 6h (G). N=3-4. Mean±STDEV. Statistical analysis performed with a two-way ANOVA with Tukey’s multiple comparison test. **p≤0.01, ****p≤0.0001.

**Supplemental Figure 12.**
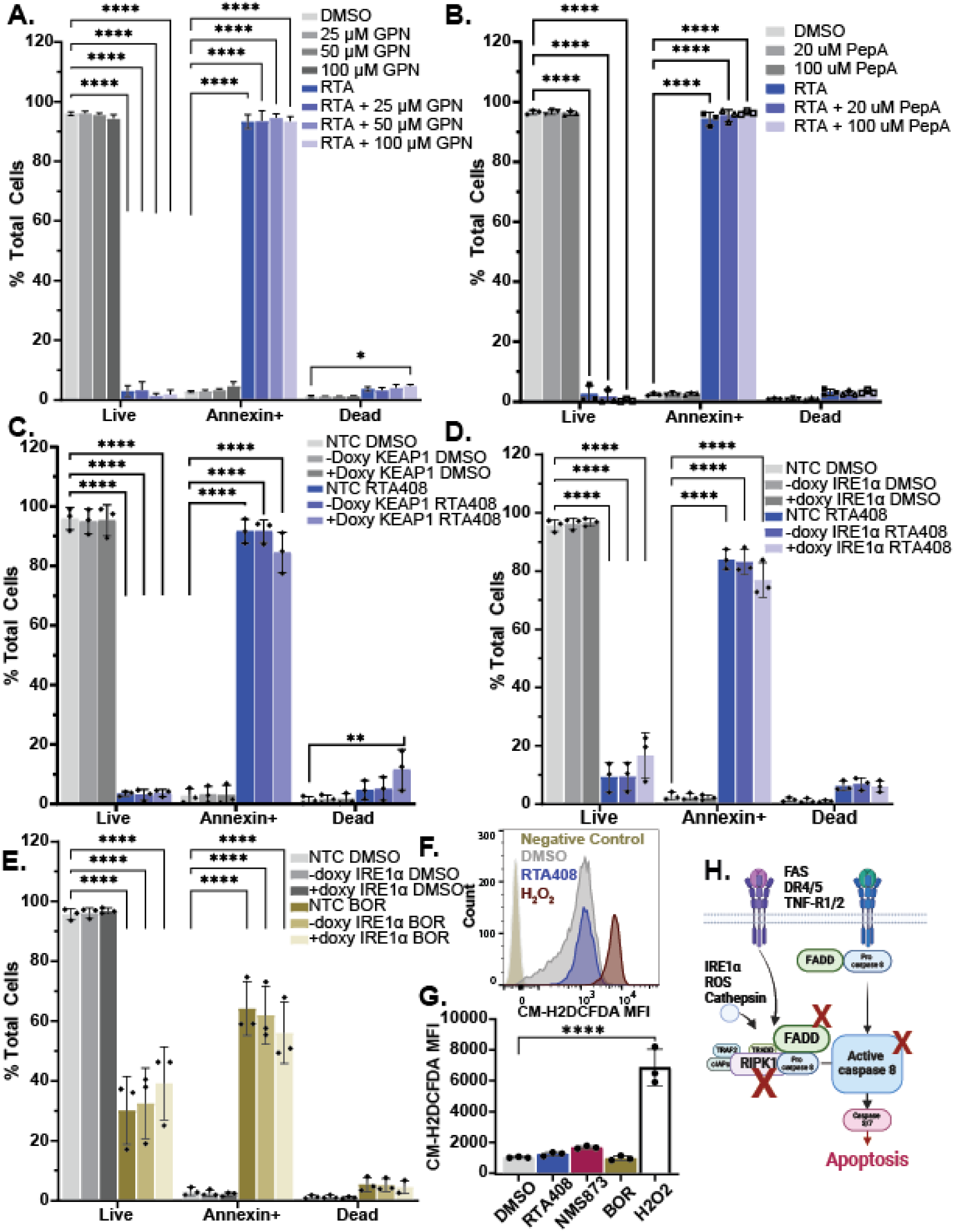
Evaluation of Intracellular Pathways Implicated with Caspase 8 Activation. A-B. Flow cytometry quantitation of live (AnnexinV-DAPI-), Annexin+ (AnnexinV+DAPI-) or dead (AnnexinV+DAPI+) in MM.1S cells treated with Gly-Phe-β-Naphthylamide (GPN-A) or Pepstatin A (PepA-B) in combination with DMSO or 1 µM RTA408 for 4h. C-E. Flow cytometry quantitation of Annexin V staining in MM.1S cells transduced with non-targeting control, or doxycycline inducible KEAP1 treated with DMSO or 1 µM RTA408 for 4h (C) or IRE1α treated DMSO, 1 µM RTA408 for 4h (D) or 120 nM BOR (E) for 6h. F-G. Representative histogram (F) and flow cytometry quantitation (G) of CM-H2DCFDA mean fluorescence intensity (MFI) in live MM.1S cells. H. Graphical summary of caspase 8 dependent pro-apoptotic signaling implicated with ERAD inhibition. N=3. Mean±STDEV statistical analysis performed with a two-way ANOVA with Tukey’s multiple comparison test. ****p≤0.0001.

**Supplement 13:**
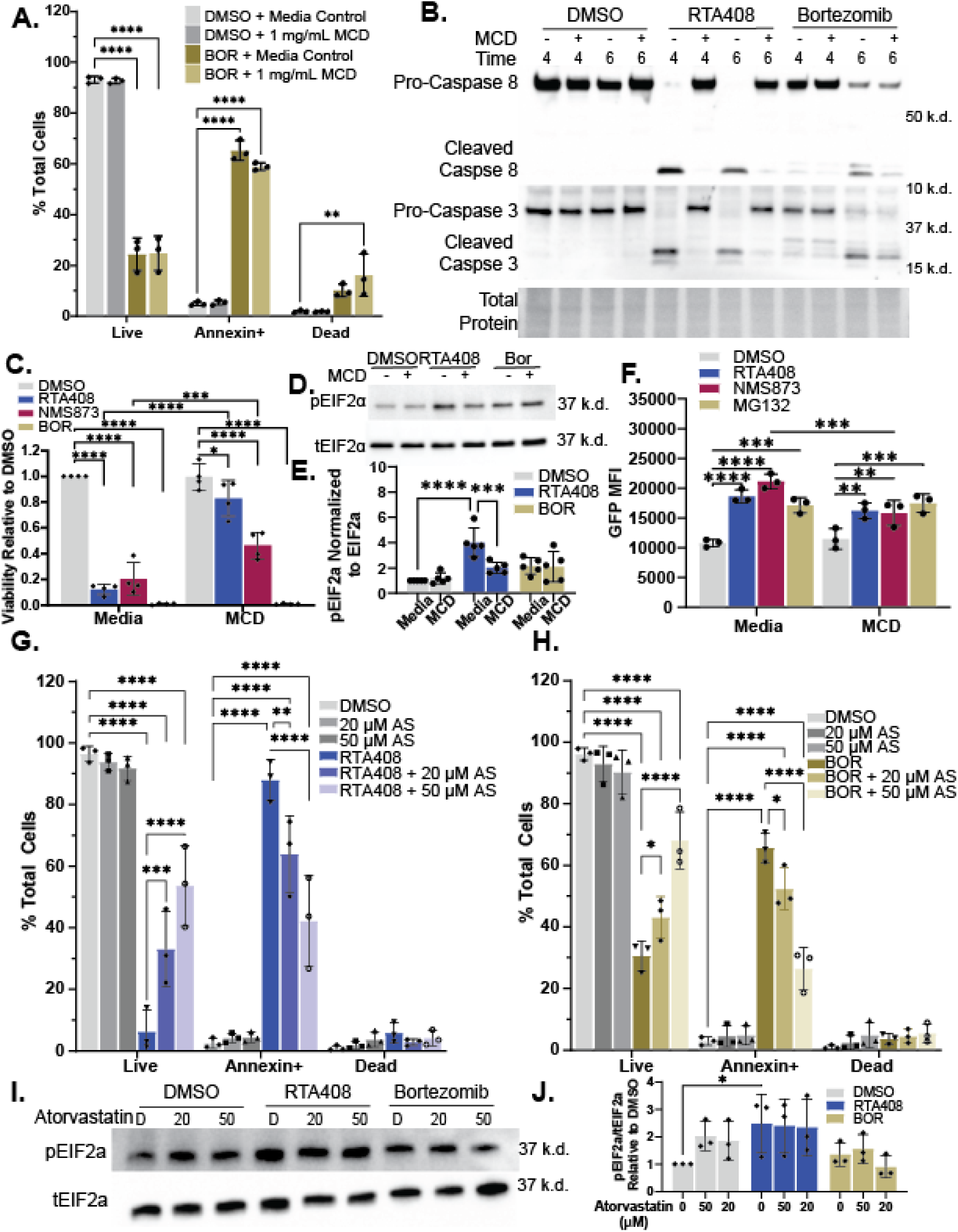
Lipid Raft Dependent Activation of Caspase 8 in MM.1S. A. Flow cytometry quantitation of Annexin V staining in MM.1S treated with 1 mg/mL MCD or media control and DMSO or 120 nM BOR for 6h. B. Representative immunoblot of caspase 8 and 3 (cleaved and pro-forms) and total protein in MM.1s treated with DMSO, 1 µM RTA408 or 120 nM BOR and media control or 1 mg/mL MCD for 4-6h. C. MM cell line viability measured by CellTiter-Glo® following treatment with 1 µM RTA408, 10 µM NMS873 or 120 nM BOR and media control or 1 mg/mL MCD for 12h. D-E. Representative immunoblot (D) and relative quantitation (E) of phospho-EIF2α or total EIF2 α with DMSO, 1 µM RTA408 or 120 nM BOR and media control or 1 mg/mL MCD for 2h. F. Steady state degradation of NHK-GFP in K562 with 10 µM RTA408, NMS873 or MG132 and media control or 1 mg/mL MCD for 4h. G-H. Flow cytometry quantitation of Annexin V staining in MM.1S cells treated with DMSO or 20-50 µM atorvastatin (AS) and DMSO or 1 µM RTA408 for 4h (G) or 120 nM BOR for 6h (H). I-J. Representative immunoblot (I) and relative quantitation (J) of phospho-EIF2α or total EIF2α with DMSO, 1 µM RTA408 or 120 nM BOR and DMSO or 20-50 µM atorvastatin (AS) for 2h. N=3-5. Mean±STDEV. Statistical analysis performed with a two-way ANOVA with Dunnett’s multiple comparison test (E) or Tukey’s multiple comparison tests. *p≤0.05, **p≤0.01, *** p≤0.001, p****p≤0.0001.

**Supplemental Figure 14:**
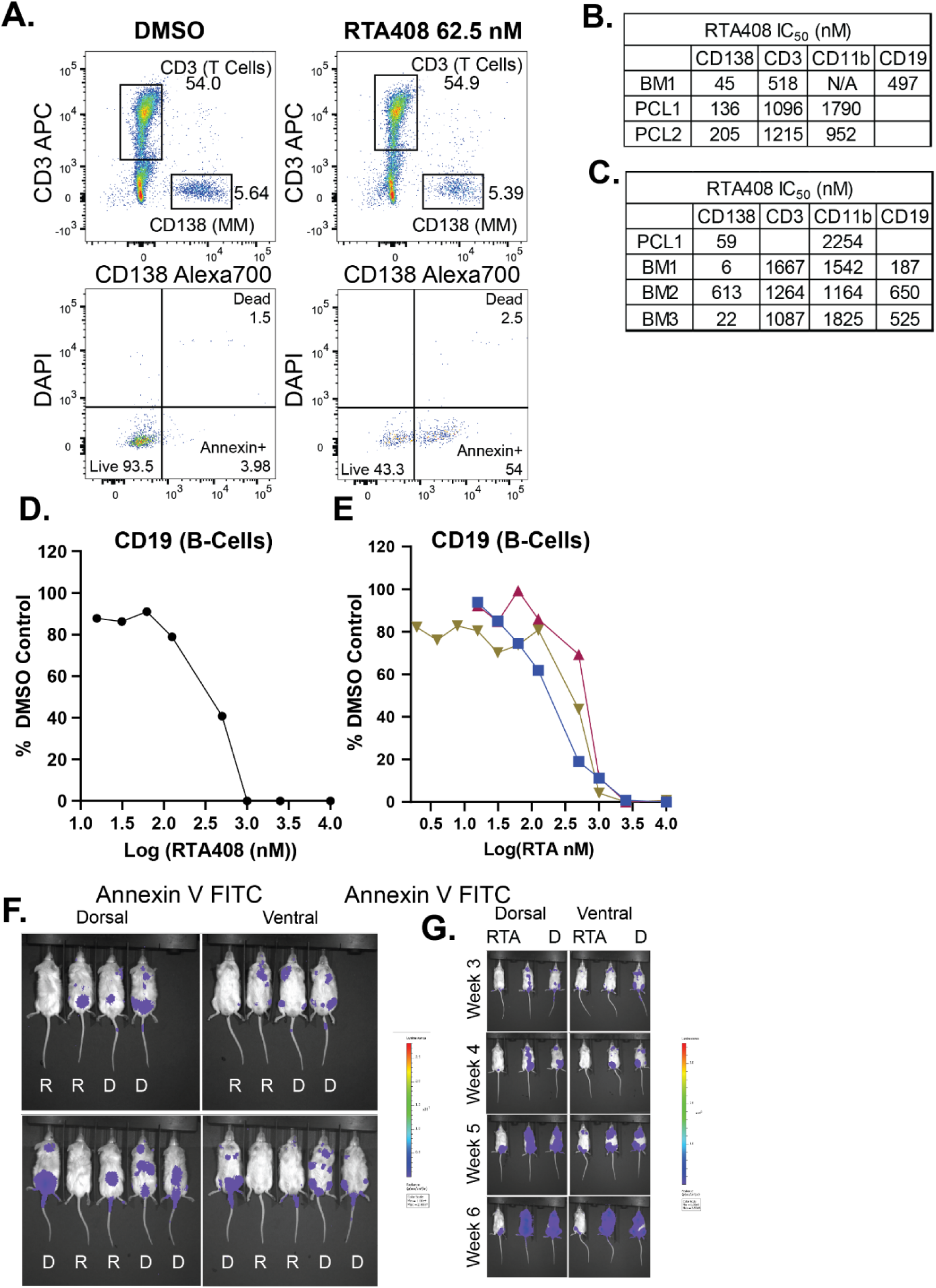
RTA408 Cytotoxicity in Primary Cells and In Vivo. A. Representative flow cytometry gating of CD138+,CD3+ and Live (Annexin-DAPI-), Annexin+ (Annexin+DAPI-), and Dead (DAPI+) cells. B-C. Calculation of IC50 for cytotoxicity in different cell populations R/R (B) and newly diagnosed (C) samples. D-E. Flow cytometry analysis of Annexin-DAPI-CD19 cells following RTA408 treatment for 60h in R/R (D) and 36h (E) in newly diagnosed MM from primary bone marrow (BM) or peripheral blood (PCL) cells from patients with relapsed refractory MM. F. Week 3, pretreatment bioluminescent imaging in RPMI8226 NSG mice. G. Bioluminescent imaging from a pilot cohort of RPMI8226 NSG mice; left treated with RTA408 5 mg/kg; right DMSO control. Center mouse not included in treatment schema.

## References

1. Christianson, J.C., and Carvalho, P. (2022). Order through destruction: how ER-associated protein degradation contributes to organelle homeostasis. EMBO J 41, e109845. 10.15252/EMBJ.2021109845.

2. Wang, Q., Mora-Jensen, H., Weniger, M.A., Perez-Galan, P., Wolford, C., Hai, T., Ron, D., Chen, W., Trenkle, W., Wiestner, A., et al. (2009). ERAD inhibitors integrate ER stress with an epigenetic mechanism to activate BH3-only protein NOXA in cancer cells. Proc Natl Acad Sci U S A 106, 2200. 10.1073/PNAS.0807611106.

3. Nishimura, N., Radwan, M.O., Amano, M., Endo, S., Fujii, E., Hayashi, H., Ueno, S., Ueno, N., Tatetsu, H., Hata, H., et al. (2019). Novel p97/VCP inhibitor induces endoplasmic reticulum stress and apoptosis in both bortezomib-sensitive and -resistant multiple myeloma cells. Cancer Sci 110, 3275. 10.1111/CAS.14154.

4. Le Moigne, R., Aftab, B.T., Djakovic, S., Dhimolea, E., Valle, E., Murnane, M., King, E.M., Soriano, F., Menon, M.K., Wu, Z.Y., et al. (2017). The p97 inhibitor CB-5083 is a unique disrupter of protein homeostasis in models of multiple myeloma. Mol Cancer Ther 16, 2375–2386. 10.1158/1535-7163.MCT-17-0233.

5. Yagishita, N., Aratani, S., Leach, C., Amano, T., Yamano, Y., Nakatani, K., Nishioka, K., and Nakajima, T. (2012). RING-finger type E3 ubiquitin ligase inhibitors as novel candidates for the treatment of rheumatoid arthritis. Int J Mol Med 30, 1281. 10.3892/IJMM.2012.1129.

6. Leinonen, H., Cheng, C., Pitkänen, M., Sander, C.L., Zhang, J., Saeid, S., Turunen, T., Shmara, A., Weiss, L., Ta, L., et al. (2021). A p97/valosin-containing protein inhibitor drug CB-5083 has a potent but reversible off-target effect on phosphodiesterase-6. Journal of Pharmacology and Experimental Therapeutics 378, 31–41. 10.1124/JPET.120.000486/-/DCSUPPLEMENTAL.

7. Aronson, L.I., and Davies, F.E. (2012). DangER: protein ovERload. Targeting protein degradation to treat myeloma. Haematologica 97, 1119. 10.3324/HAEMATOL.2012.064923.

8. Kumar, S.K., Rajkumar, V., Kyle, R.A., Van Duin, M., Sonneveld, P., Mateos, M.V., Gay, F., and Anderson, K.C. (2017). Multiple myeloma. Nat Rev Dis Primers 3, 1–20. 10.1038/nrdp.2017.46.

9. Obeng, E.A., Carlson, L.M., Gutman, D.M., Harrington, W.J., Lee, K.P., and Boise, L.H. (2006). Proteasome inhibitors induce a terminal unfolded protein response in multiple myeloma cells. Blood 107, 4907–4916. 10.1182/BLOOD-2005-08-3531.

10. Calame, K.L., Lin, K.I., and Tunyaplin, C. (2003). Regulatory mechanisms that determine the development and function of plasma cells. Annu Rev Immunol 21, 205–230. 10.1146/ANNUREV.IMMUNOL.21.120601.141138.

11. Mitsiades, N., Mitsiades, C.S., Poulaki, V., Chauhan, D., Fanourakis, G., Gu, X., Bailey, C., Joseph, M., Libermann, T.A., Treon, S.P., et al. (2002). Molecular sequelae of proteasome inhibition in human multiple myeloma cells. Proc Natl Acad Sci U S A 99, 14374. 10.1073/PNAS.202445099.

12. de Matos Simoes, R., Shirasaki, R., Downey-Kopyscinski, S.L., Matthews, G.M., Barwick, B.G., Gupta, V.A., Dupéré-Richer, D., Yamano, S., Hu, Y., Sheffer, M., et al. (2023). Genome-Scale Functional Genomics Identify Genes Preferentially Essential for Multiple Myeloma Cells compared to other Neoplasias. Nat Cancer 4, 754. 10.1038/S43018-023-00550-X.

13. Joseph, N.S., Kaufman, J.L., Dhodapkar, M. V., Hofmeister, C.C., Almaula, D.K., Heffner, L.T., Gupta, V.A., Boise, L.H., Lonial, S., and Nooka, A.K. (2020). Long-Term Follow-Up Results of Lenalidomide, Bortezomib, and Dexamethasone Induction Therapy and Risk-Adapted Maintenance Approach in Newly Diagnosed Multiple Myeloma. Journal of Clinical Oncology 38, 1928. 10.1200/JCO.19.02515.

14. Abramson, H.N. (2018). The Multiple Myeloma Drug Pipeline—2018: A Review of Small Molecules and Their Therapeutic Targets. Clin Lymphoma Myeloma Leuk 18, 611–627. 10.1016/J.CLML.2018.06.015.

15. Nandi, P., DeVore, K., Wang, F., Li, S., Walker, J.D., Truong, T.T., LaPorte, M.G., Wipf, P., Schlager, H., McCleerey, J., et al. (2024). Mechanism of allosteric inhibition of human p97/VCP ATPase and its disease mutant by triazole inhibitors. Communications Chemistry 2024 7:1 7, 1–14. 10.1038/s42004-024-01267-3.

16. Lynch, D.R., Farmer, J., Hauser, L., Blair, I.A., Wang, Q.Q., Mesaros, C., Snyder, N., Boesch, S., Chin, M., Delatycki, M.B., et al. (2019). Safety, pharmacodynamics, and potential benefit of omaveloxolone in Friedreich ataxia. Ann Clin Transl Neurol 6, 15–26. 10.1002/acn3.660.

17. Lynch, D.R., Chin, M.P., Boesch, S., Delatycki, M.B., Giunti, P., Goldsberry, A., Hoyle, J.C., Mariotti, C., Mathews, K.D., Nachbauer, W., et al. (2023). Efficacy of Omaveloxolone in Friedreich’s Ataxia: Delayed-Start Analysis of the MOXIe Extension. Movement Disorders 38, 313–320. 10.1002/MDS.29286.

18. Leto, D.E., Morgens, D.W., Zhang, L., Walczak, C.P., Elias, J.E., Bassik, M.C., and Kopito, R.R. (2019). Genome-wide CRISPR analysis identifies substrate-specific conjugation modules in ER-associated degradation. Mol Cell 73, 377. 10.1016/J.MOLCEL.2018.11.015.

19. Jacob, R.T., Larsen, M.J., Larsen, S.D., Kirchhoff, P.D., Sherman, D.H., and Neubig, R.R. (2012). MScreen: an integrated compound management and high-throughput screening data storage and analysis system. J Biomol Screen 17, 1080–1087. 10.1177/1087057112450186.

20. Yanamandra, N., Colaco, N.M., Parquet, N.A., Buzzeo, R.W., Boulware, D., Wright, G., Perez, L.E., Dalton, W.S., and Beaupre, D.M. (2006). Tipifarnib and bortezomib are synergistic and overcome cell adhesion-mediated drug resistance in multiple myeloma and acute myeloid leukemia. Clin Cancer Res 12, 591–599. 10.1158/1078-0432.CCR-05-1792.

21. Ria, R., Melaccio, A., Racanelli, V., and Vacca, A. (2020). Anti-VEGF Drugs in the Treatment of Multiple Myeloma Patients. J Clin Med 9, 1765. 10.3390/JCM9061765.

22. Rossi, M., Teresa Di Martino, M., Morelli, E., Leotta, M., Rizzo, A., Grimaldi, A., Misso, G., Tassone, P., and Caraglia, M. (2012). Molecular targets for the treatment of multiple myeloma. Curr Cancer Drug Targets 12, 757–767. 10.2174/156800912802429300.

23. Günther, A., Baumann, P., Burger, R., Kellner, C., Klapper, W., Schmidmaier, R., and Gramatzki, M. (2015). Activity of everolimus (RAD001) in relapsed and/or refractory multiple myeloma: a phase I study. Haematologica 100, 541–547. 10.3324/HAEMATOL.2014.116269.

24. Li, J., Liu, Z., Li, Y., Jing, Q., Wang, H., Liu, H., Chen, J., Feng, J., Shao, Q., and Fu, R. (2019). Everolimus shows synergistic antimyeloma effects with bortezomib via the AKT/mTOR pathway. J Investig Med 67, 39–47. 10.1136/JIM-2018-000780.

25. Xu, B., Li, J., Xu, D., and Ran, Q. (2023). PLK4 inhibitor plus bortezomib exhibits a synergistic effect on treating multiple myeloma via inactivating PI3K/AKT signaling. Ir J Med Sci 192, 561–567. 10.1007/S11845-022-03007-9.

26. Consortium, T.U., Bateman, A., Martin, M.-J., Orchard, S., Magrane, M., Adesina, A., Ahmad, S., Bowler-Barnett, E.H., Bye-A-Jee, H., Carpentier, D., et al. (2025). UniProt: the Universal Protein Knowledgebase in 2025. Nucleic Acids Res 53, D609–D617. 10.1093/NAR/GKAE1010.

27. Jovanović, K.K., Roche-Lestienne, C., Ghobrial, I.M., Facon, T., Quesnel, B., and Manier, S. (2018). Targeting MYC in multiple myeloma. Leukemia 2018 32:6 32, 1295–1306. 10.1038/s41375-018-0036-x.

28. Hideshima, T., Ikeda, H., Chauhan, D., Okawa, Y., Raje, N., Podar, K., Mitsiades, C., Munshi, N.C., Richardson, P.G., Carrasco, R.D., et al. (2009). Bortezomib induces canonical nuclear factor-κB activation in multiple myeloma cells. Blood 114, 1046–1052. 10.1182/BLOOD-2009-01-199604.

29. Pakjoo, M., Ahmadi, S.E., Zahedi, M., Jaafari, N., Khademi, R., Amini, A., and Safa, M. (2024). Interplay between proteasome inhibitors and NF-κB pathway in leukemia and lymphoma: a comprehensive review on challenges ahead of proteasome inhibitors. Cell Communication and Signaling 2024 22:1 22, 1–28. 10.1186/S12964-023-01433-5.

30. Ohtsuki, T., Yawata, Y., Wada, H., Sugihara, T., Mori, M., and Namba, M. (1989). Two human myeloma cell lines, amylase-producing KMS-12-PE and amylase-non-producing KMS-12-BM, were established from a patient, having the same chromosome marker, t(11;14)(q13;q32). Br J Haematol 73, 199–204. 10.1111/J.1365-2141.1989.TB00252.X.

31. Zhou, L. (2022). Caspase-8: Friend or Foe in Bortezomib/Lenalidomide-Based Therapy for Myeloma. Front Oncol 12. 10.3389/FONC.2022.861709.

32. Chen, W., Sun, M., Sun, Y., Yang, Q., Gao, H., Li, L., Fu, R., and Dong, N. (2024). Proteasome Inhibition Induces Apoptosis Through Simultaneous Inactivation of MCL-1/BCL-XL by NOXA Independent of CHOP and JNK Pathways. Toxicology, 153906. 10.1016/J.TOX.2024.153906.

33. Mitsiades, N., Mitsiades, C.S., Poulaki, V., Chauhan, D., Fanourakis, G., Gu, X., Bailey, C., Joseph, M., Libermann, T.A., Treon, S.P., et al. (2002). Molecular sequelae of proteasome inhibition in human multiple myeloma cells. Proc Natl Acad Sci U S A 99, 14374–14379. 10.1073/PNAS.202445099/SUPPL_FILE/4450FIG6.JPG.

34. Ali, M., and Mocarski, E.S. (2018). Proteasome inhibition blocks necroptosis by attenuating death complex aggregation. Cell Death & Disease 2018 9:3 9, 1–12. 10.1038/s41419-018-0371-x.

35. Hellwig, C.T., Delgado, M.E., Skoko, J., Dyck, L., Hanna, C., Wentges, A., Langlais, C., Hagenlocher, C., Mack, A., Dinsdale, D., et al. (2021). Proteasome inhibition triggers the formation of TRAIL receptor 2 platforms for caspase-8 activation that accumulate in the cytosol. Cell Death & Differentiation 2021 29:1 29, 147–155. 10.1038/s41418-021-00843-7.

36. Tian, X., Srinivasan, P.R., Tajiknia, V., Sanchez Sevilla Uruchurtu, A.F., Seyhan, A.A., Carneiro, B.A., De La Cruz, A., Pinho-Schwermann, M., George, A., Zhao, S., et al. (2024). Targeting apoptotic pathways for cancer therapy. J Clin Invest 134, e179570. 10.1172/JCI179570.

37. Stanger, B.Z., Leder, P., Lee, T.H., Kim, E., and Seed, B. (1995). RIP: a novel protein containing a death domain that interacts with Fas/APO-1 (CD95) in yeast and causes cell death. Cell 81, 513–523. 10.1016/0092-8674(95)90072-1.

38. Hsu, H., Xiong, J., and Goeddel, D. V. (1995). The TNF receptor 1-associated protein TRADD signals cell death and NF-kappa B activation. Cell 81, 495–504. 10.1016/0092-8674(95)90070-5.

39. Ju, E., Park, K.A., Shen, H.M., and Hur, G.M. (2022). The resurrection of RIP kinase 1 as an early cell death checkpoint regulator—a potential target for therapy in the necroptosis era. Experimental & Molecular Medicine 2022 54:9 54, 1401–1411. 10.1038/s12276-022-00847-4.

40. Rauert, H., Stühmer, T., Bargou, R., Wajant, H., and Siegmund, D. (2011). TNFR1 and TNFR2 regulate the extrinsic apoptotic pathway in myeloma cells by multiple mechanisms. Cell Death Dis 2, e194. 10.1038/CDDIS.2011.78.

41. Ren, J., Zhang, X., Liu, X., Fang, C., Jiang, S., June, C.H., and Zhao, Y. (2017). A versatile system for rapid multiplex genome-edited CAR T cell generation. Oncotarget 8, 17002. 10.18632/ONCOTARGET.15218.

42. Shemorry, A., Harnoss, J.M., Guttman, O., Marsters, S.A., Kőműves, L.G., Lawrence, D.A., and Ashkenazi, A. (2019). Caspase-mediated cleavage of IRE1 controls apoptotic cell commitment during endoplasmic reticulum stress. Elife 8. 10.7554/eLife.47084.

43. Sano, R., and Reed, J.C. (2013). ER stress-induced cell death mechanisms. Biochimica et Biophysica Acta (BBA) - Molecular Cell Research 1833, 3460–3470. 10.1016/J.BBAMCR.2013.06.028.

44. Redza-Dutordoir, M., and Averill-Bates, D.A. (2016). Activation of apoptosis signalling pathways by reactive oxygen species. Biochimica et Biophysica Acta (BBA) - Molecular Cell Research 1863, 2977–2992. 10.1016/J.BBAMCR.2016.09.012.

45. Conus, S., Pop, C., Snipas, S.J., Salvesen, G.S., and Simon, H.U. (2012). Cathepsin D Primes Caspase-8 Activation by Multiple Intra-chain Proteolysis. J Biol Chem 287, 21142. 10.1074/JBC.M111.306399.

46. Baumgartner, H.K., Gerasimenko, J. V., Thorne, C., Ashurst, L.H., Barrow, S.L., Chvanov, M.A., Gillies, S., Criddle, D.N., Tepikin, A. V., Petersen, O.H., et al. (2007). Caspase-8-mediated apoptosis induced by oxidative stress is independent of the intrinsic pathway and dependent on cathepsins. Am J Physiol Gastrointest Liver Physiol 293, 296–307. 10.1152/AJPGI.00103.2007/ASSET/IMAGES/LARGE/ZH30070748130010.JPEG.

47. Estornes, Y., Aguileta, M.A., Dubuisson, C., De Keyser, J., Goossens, V., Kersse, K., Samali, A., Vandenabeele, P., and Bertrand, M.J.M. (2014). RIPK1 promotes death receptor-independent caspase-8-mediated apoptosis under unresolved ER stress conditions. Cell Death & Disease 2014 5:12 5, e1555–e1555. 10.1038/cddis.2014.523.

48. Mollinedo, F., De La Iglesia-Vicente, J., Gajate, C., Estella-Hermoso De Mendoza, A., Villa-Pulgarin, J.A., Campanero, M.A., and Blanco-Prieto, M.J. (2010). Lipid raft-targeted therapy in multiple myeloma. Oncogene 2010 29:26 29, 3748–3757. 10.1038/onc.2010.131.

49. Rozemuller, H., Van Der Spek, E., Bogers-Boer, L.H., Zwart, M.C., Verweij, V., Emmelot, M., Groen, R.W., Spaapen, R., Bloem, A.C., Lokhorst, H.M., et al. (2008). A bioluminescence imaging based in vivo model for preclinical testing of novel cellular immunotherapy strategies to improve the graft-versus-myeloma effect. Haematologica 93, 1049–1057. 10.3324/haematol.12349.

50. Mehdi, S.H., Nafees, S., Mehdi, S.J., Morris, C.A., Mashouri, L., and Yoon, D. (2021). Animal Models of Multiple Myeloma Bone Disease. Front Genet 12, 640954. 10.3389/FGENE.2021.640954/BIBTEX.

51. Bhattacharya, A., and Qi, L. (2019). ER-associated degradation in health and disease – from substrate to organism. J Cell Sci 132. 10.1242/JCS.232850.

52. Dayalan Naidu, S., Muramatsu, A., Saito, R., Asami, S., Honda, T., Hosoya, T., Itoh, K., Yamamoto, M., Suzuki, T., and Dinkova-Kostova, A.T. (2018). C151 in KEAP1 is the main cysteine sensor for the cyanoenone class of NRF2 activators, irrespective of molecular size or shape. Sci Rep 8, 8037. 10.1038/S41598-018-26269-9.

53. Kim, T.Y., Kim, E., Yoon, S.K., and Yoon, J.B. (2008). Herp enhances ER-associated protein degradation by recruiting ubiquilins. Biochem Biophys Res Commun 369, 741–746. 10.1016/J.BBRC.2008.02.086.

54. Huang, C.H., Chu, Y.R., Ye, Y., and Chen, X. (2014). Role of HERP and a HERP-related protein in HRD1-dependent protein degradation at the endoplasmic reticulum. Journal of Biological Chemistry 289, 4444–4454. 10.1074/jbc.M113.519561.

55. Patiño-Escobar, B., Talbot, A., and Wiita, A.P. (2023). Overcoming proteasome inhibitor resistance in the immunotherapy era. Trends Pharmacol Sci 44, 507–518. 10.1016/j.tips.2023.05.006.

56. Jovanović, K.K., Roche-Lestienne, C., Ghobrial, I.M., Facon, T., Quesnel, B., and Manier, S. (2018). Targeting MYC in multiple myeloma. Leukemia 2018 32:6 32, 1295–1306. 10.1038/s41375-018-0036-x.

57. Bird, S., and Pawlyn, C. (2023). IMiD resistance in multiple myeloma: current understanding of the underpinning biology and clinical impact. Blood 142, 131–140. 10.1182/BLOOD.2023019637.

58. Walter, P., and Ron, D. (2011). The Unfolded Protein Response: From Stress Pathway to Homeostatic Regulation. Science (1979) 334, 1081–1086. 10.1126/SCIENCE.1209038.

59. Auner, H.W., Moody, A.M., Ward, T.H., Kraus, M., Milan, E., May, P., Chaidos, A., Driessen, C., Cenci, S., Dazzi, F., et al. (2013). Combined Inhibition of p97 and the Proteasome Causes Lethal Disruption of the Secretory Apparatus in Multiple Myeloma Cells. PLoS One 8. 10.1371/JOURNAL.PONE.0074415.

60. Rauert, H., Stühmer, T., Bargou, R., Wajant, H., and Siegmund, D. (2011). TNFR1 and TNFR2 regulate the extrinsic apoptotic pathway in myeloma cells by multiple mechanisms. Cell Death Dis 2. 10.1038/CDDIS.2011.78.

61. Hellwig, C.T., Delgado, M.E., Skoko, J., Dyck, L., Hanna, C., Wentges, A., Langlais, C., Hagenlocher, C., Mack, A., Dinsdale, D., et al. (2021). Proteasome inhibition triggers the formation of TRAIL receptor 2 platforms for caspase-8 activation that accumulate in the cytosol. Cell Death & Differentiation 2021 29:1 29, 147–155. 10.1038/s41418-021-00843-7.

62. Stevenson, J., Huang, E.Y., and Olzmann, J.A. (2016). Endoplasmic Reticulum–Associated Degradation and Lipid Homeostasis. Annu Rev Nutr 36, 511. 10.1146/ANNUREV-NUTR-071715-051030.

63. Mollinedo, F., and Gajate, C. (2015). Lipid rafts as major platforms for signaling regulation in cancer. Adv Biol Regul 57, 130–146. 10.1016/J.JBIOR.2014.10.003.

64. Boldin, M.P., Mett, I.L., Varfolomeev, E.E., Chumakov, I., Shemer-Avni, Y., Camonis, J.H., and Wallach, D. (1995). Self-association of the “Death Domains” of the p55 Tumor Necrosis Factor (TNF) Receptor and Fas/APO1 Prompts Signaling for TNF and Fas/APO1 Effects. Journal of Biological Chemistry 270, 387–391. 10.1074/JBC.270.1.387.

65. Lincoln, J.E., Boling, M., Parikh, A.N., Yeh, Y., Gilchrist, D.G., and Morse, L.S. (2006). Fas signaling induces raft coalescence that is blocked by cholesterol depletion in human RPE cells undergoing apoptosis. Invest Ophthalmol Vis Sci 47, 2172–2178. 10.1167/IOVS.05-1167.

66. Ji, Y., Kim, H., Yang, L., Sha, H., Roman, C.A., Long, Q., and Qi, L. (2016). The Sel1L-Hrd1 endoplasmic reticulum-associated degradation complex manages a key checkpoint in B cell development. Cell Rep 16, 2630. 10.1016/J.CELREP.2016.08.003.

67. Manganelli, V., Longo, A., Mattei, V., Recalchi, S., Riitano, G., Caissutti, D., Capozzi, A., Sorice, M., Misasi, R., and Garofalo, T. (2021). Role of ERLINs in the Control of Cell Fate through Lipid Rafts. Cells 10, 2408. 10.3390/CELLS10092408.

68. Pearce, M.M.P., Wormer, D.B., Wilkens, S., and Wojcikiewicz, R.J.H. (2009). An Endoplasmic Reticulum (ER) Membrane Complex Composed of SPFH1 and SPFH2 Mediates the ER-associated Degradation of Inositol 1,4,5-Trisphosphate Receptors. J Biol Chem 284, 10433. 10.1074/JBC.M809801200.

69. Rückrich, T., Kraus, M., Gogel, J., Beck, A., Ovaa, H., Verdoes, M., Overkleeft, H.S., Kalbacher, H., and Driessen, C. (2009). Characterization of the ubiquitin-proteasome system in bortezomib-adapted cells. Leukemia 23, 1098–1105. 10.1038/LEU.2009.8.

70. Zaal, E.A., Wu, W., Jansen, G., Zweegman, S., Cloos, J., and Berkers, C.R. (2017). Bortezomib resistance in multiple myeloma is associated with increased serine synthesis. Cancer Metab 5, 7. 10.1186/S40170-017-0169-9.

71. McAlister, G.C., Nusinow, D.P., Jedrychowski, M.P., Wühr, M., Huttlin, E.L., Erickson, B.K., Rad, R., Haas, W., and Gygi, S.P. (2014). MultiNotch MS3 enables accurate, sensitive, and multiplexed detection of differential expression across cancer cell line proteomes. Anal Chem 86, 7150–7158. 10.1021/AC502040V.

72. Aubrey, B.J., Kelly, G.L., Kueh, A.J., Brennan, M.S., O’Connor, L., Milla, L., Wilcox, S., Tai, L., Strasser, A., and Herold, M.J. (2015). An inducible lentiviral guide RNA platform enables the identification of tumor-essential genes and tumor-promoting mutations in vivo. Cell Rep 10, 1422–1432. 10.1016/J.CELREP.2015.02.002.

73. Zhang, J.H., Chung, T.D.Y., and Oldenburg, K.R. (1999). A Simple Statistical Parameter for Use in Evaluation and Validation of High Throughput Screening Assays. J Biomol Screen 4, 67–73. 10.1177/108705719900400206.

74. Perez-Riverol, Y., Bandla, C., Kundu, D.J., Kamatchinathan, S., Bai, J., Hewapathirana, S., John, N.S., Prakash, A., Walzer, M., Wang, S., et al. (2024). The PRIDE database at 20 years: 2025 update. Nucleic Acids Res 53, D543. 10.1093/NAR/GKAE1011.

